# Improved architectures for flexible DNA production using retrons across kingdoms of life

**DOI:** 10.1101/2021.03.26.437017

**Authors:** Santiago C. Lopez, Kate D. Crawford, Santi Bhattarai-Kline, Seth L. Shipman

**Author notes:** Equal Contributions (Santiago C. Lopez, Kate D. Crawford).

## Abstract

Exogenous DNA is a critical molecular tool for biology. This is particularly true for gene editing, where exogenous DNA can be used as a template to introduce precise changes to the sequence of a cell’s genome. This DNA is typically synthesized or assembled *in vitro* and then delivered to target cells. However, delivery can be inefficient, and low abundance of template DNA may be one reason that precise editing typically occurs at a low rate. It has recently been shown that producing DNA inside cells can using reverse transcriptases can increase the efficiency of genome editing. One tool to produce that DNA is a retron, a bacterial retroelement that has an endogenous role in phage defense. However, little effort has been directed at optimizing the retron for production of designed sequences when used as a component of biotechnology. Here, we identify modifications to the retron non-coding RNA that result in more abundant reverse transcribed DNA. We also test architectures of the retron operon that enable efficient reverse transcription across kingdoms of life from bacteria, to yeast, to cultured human cells. We find that gains in DNA production using modified retrons are portable from prokaryotic to eukaryotic cells. Finally, we demonstrate that increased production of RT-DNA results in more efficient genome editing in both prokaryotic and eukaryotic cells. These experiments provide a general framework for production of DNA using a retron for biotechnological applications.

## INTRODUCTION

Exogenous DNA, which does not match the genome of the cell where it is harbored, is a fundamental tool of modern cell and molecular biology. This DNA can serve as a template to modify a cell’s genome, subtly altering existing genes, or even inserting wholly new genetic material that adds function or marks a cellular event such as lineage. Exogenous DNA for these uses is typically synthesized or assembled in a tube, then physically delivered to the cells that will be altered. However, it remains an incredible challenge to deliver exogenous DNA to cells in universally high abundance and without substantial variation between recipients^1^. These technical challenges likely contribute to low rates of precise editing as well as unintended editing that occurs in the absence of template DNA^2–4^. Effort has been made to bias cells toward template-based editing by manipulating the proteins involved in DNA repair or tethering DNA templates to other editing materials to increase their local concentration^5^. However, a simpler approach may be to eliminate DNA delivery problems by producing the DNA inside the cell.

In recent years, it has been shown that retroelements can be used to produce DNA for genome editing within cells by reverse transcription^6–9^. This reverse transcribed DNA (RT-DNA) is produced in cells from a universally delivered cassette in high abundance such that it overcomes inefficiencies and increases rates of genome editing. One retroelement class that has been useful in this regard are bacterial retrons^6, 8, 9^, which are elements involved in phage defense^10–13^. Retrons are attractive as a biotechnology due to their compact size, tightly defined sites of RT initiation and termination, lack of known host factor requirements, and lack of transposable elements. Indeed, retron-generated RT-DNA has demonstrated utility in bacterial^6, 9^ and eukaryotic^8^ genome editing.

Despite the potential of the retron as component of molecular biotechnology, it has so far been modified only as much as is necessary to show that it can produce an editing template. Given that the advantage of the retroelement approach is the increased cellular abundance of RT-DNA, we asked whether we could identify retron modifications that would yield even more abundant RT-DNA and increase in editing efficiency. Further, most work with retrons has been carried out in bacteria, with only one functional demonstration in yeast^8^, and only a brief description of reverse transcription in mammalian cells (NIH3T3 mouse cells)^14^. Therefore, we wanted to compare a more flexible architecture for retron expression across kingdoms of life, to see if there is a universal framework for RT-DNA production.

Here, we used variant libraries in *E. coli* to show that extension of complementarity in the a1/a2 region of the retron non-coding RNA (ncRNA) increases production of RT-DNA. This effect generalized across different retrons and kingdoms, from bacteria to yeast. Moreover, retron DNA production across kingdoms was possible using a universal architecture. Finally, increasing the abundance of RT-DNA in the context of genome engineering increased the rate of editing in both prokaryotic and eukaryotic cells, simultaneously showing that the template abundance is limiting for these editing applications and demonstrating a simple means of increasing efficiency.

## RESULTS

### Modifications to the retron ncRNA that affect RT-DNA production

A typical retron operon consists of a reverse transcriptase (RT), a non-coding RNA (ncRNA) that is both the primer and template for the reverse transcriptase, and one or more accessory proteins^15^ (**Fig 1a**). The RT partially reverse transcribes the ncRNA to produce a single-stranded RT-DNA with a characteristic hairpin structure, which varies in length from 48-163 bases^16^. The ncRNA can be sub-divided into a region that is reverse transcribed (msd) and a region that remains RNA in the final molecule (msr), which are partially overlapping^17–20^.

**Figure 1.**
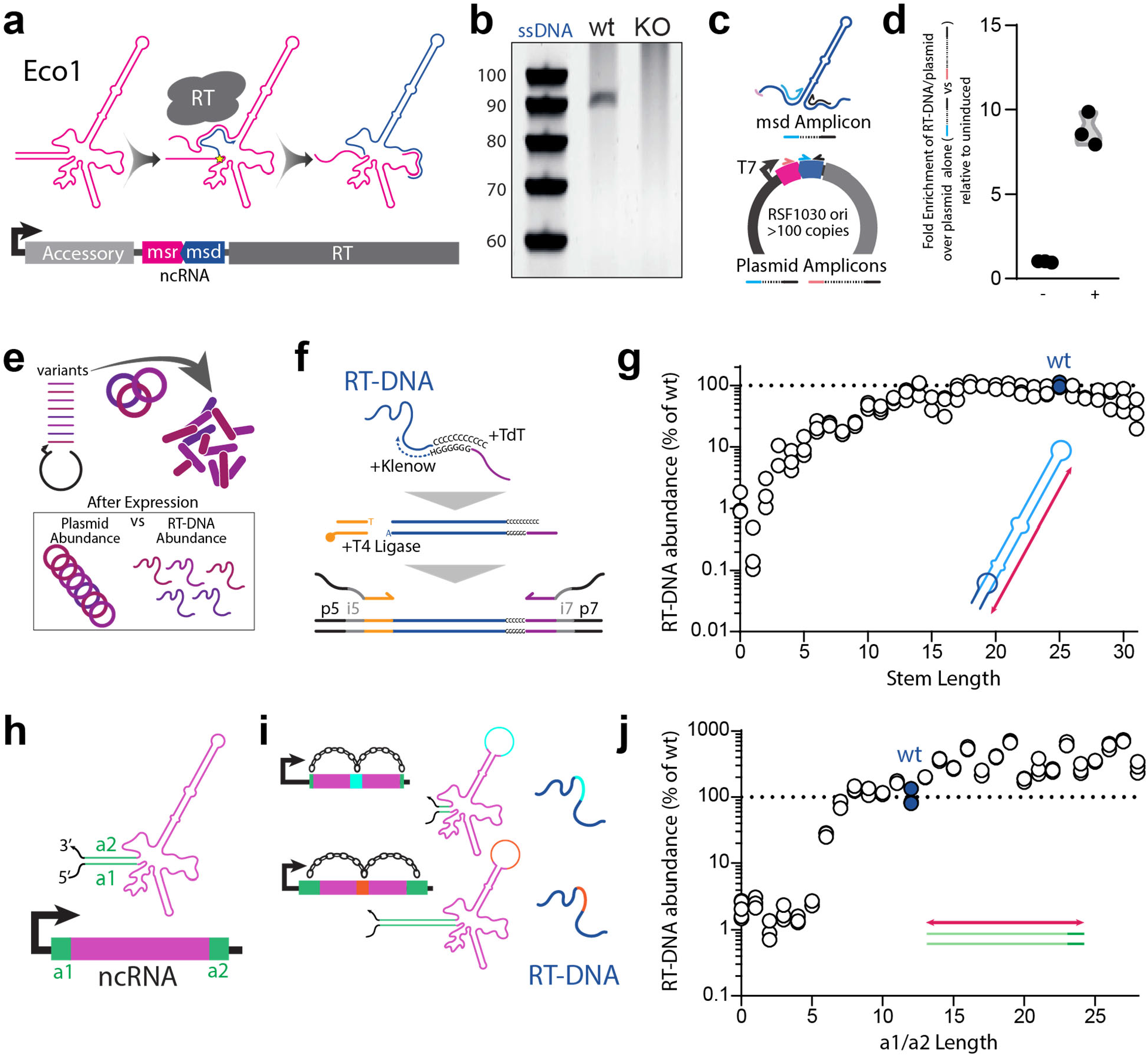
Modifications to retron ncRNA that affect RT-DNA production. **a.** Schematic of the Eco1 retron operon below, and the conversion of the ncRNA (pink) to RT-DNA (blue) above. **b.** PAGE analysis of endogenous RT-DNA produced from Eco1 in BL21-AI wild-type cells (wt) and a knockout of the retron operon (KO). **c.** qPCR analysis schematic for the RT-DNA. Blue/black primer pair will amplify using both the RT-DNA and the msd portion of the plasmid as a template. Red/black primer pair will only amplify using the plasmid as a template. **d.** Enrichment of the RT-DNA/plasmid template over the plasmid alone by qPCR, relative to the uninduced condition. Circles show each of three biological replicates. **e.** Schematic of the variant library construction and analysis. **f.** Schematic of the sequencing prep pipeline for RT-DNA. **g.** Relative RT-DNA abundance of each stem length variant as a percent of wt. Circles represent each of three biological replicates. Wt length is shown in blue along with a dashed line at 100%. **h.** Schematic illustrating the a1 and a2 regions of the retron ncRNA. **i.** Variants of the a1/a2 region are linked to a barcode in the msd loop for sequencing. **j.** Relative RT-DNA abundance of each a1/a2 length variant as a percent of wt. Circles represent each of three biological replicates. Wt length is shown in blue along with a dashed line at 100%. All statistics in Supplementary Table 1.

One of the first described retrons was found in *E. coli,* Eco1 (previously ec86)^20^. In BL21 cells, this retron is both present and active, producing RT-DNA that can be detected at the population level, which is eliminated by removing the retron operon from the genome (**Fig 1b**). In the absence of this native operon, the ncRNA and RT can be expressed from a plasmid lacking the accessory protein. Since the accessory protein is a core component of the phage-defense conferred by retrons^10–12^, this reduced system would phage defense capacity, yet continues to produce abundant RT-DNA. We quantified this RT-DNA using a relative qPCR, comparing amplification by primers against the msd region, which can use both the RT-DNA and plasmid as a template, to amplification by primers that only amplify the plasmid (**Fig 1c,d**). In *E. coli* lacking an endogenous retron, overexpression of the ncRNA and RT from a plasmid yielded an ~8-10 fold enrichment of the RT-DNA/plasmid region over the plasmid alone, which is evidence of substantial reverse transcription (**Fig 1d**).

Given that the retron utility in biotechnology relies on increasing the RT-DNA abundance in cells above what can be achieved with delivery of a synthetic template, we set out to test whether there are aspects of the ncRNA that could be modified to produce even more abundant RT-DNA. To do this, we synthesized variants of the Eco1 ncRNA and cloned them into vector for expression, with RT expressed from a separate vector. Our initial library contained variants that extended or reduced the length of the hairpin stem of the RT-DNA. This variant cloning took place in single-pot, golden gate reactions and the resulting libraries were purified and then cloned into an expression strain for analysis of RT-DNA production (**Fig 1e**). Cells harboring these library vector sets were grown overnight and then diluted and ncRNA expression was induced during growth for 5 hours.

We quantified the relative abundance of each variant plasmid in the expression strain by multiplexed Illumina sequencing before and after expression. After expression, we additionally purified RT-DNA from pools of cells harboring different retron variants by isolating cellular nucleic acids, treating that population with an RNase mixture (A/T1), and then isolating single-stranded DNA from double-stranded DNA using a commercial column-based kit. We then sequenced the RT-DNAs, comparing their relative abundance to that of their plasmid of origin to quantify the influence of different ncRNA parameters on RT-DNA production. To sequence the RT-DNA variants in this library, we used a custom sequencing pipeline to prep each RT-DNA without biasing toward any variant. This involved tailing purified RT-DNA with a string of polynucleotides using a template-independent polymerase (TdT) (**Supplemental Fig 1a**), and then generating a complementary strand via an adapter-containing, inverse anchored primer. Finally, we ligated a second adapter to this double-stranded DNA and proceed to indexing and multiplexed sequencing (**Fig 1f**).

In this first library, we modified the msd stem length from 0-31 base pairs, and found that stem length can have a large impact on RT-DNA production (**Fig 1g**). The RT tolerated modifications of the msd stem length that deviate by a small amount from the wild-type (wt) length of 25 base pairs. However, variants with stem lengths <12 and >30 produced less than half as much RT-DNA compared to the wt. Therefore, we used stem length of between 12 and 30 base pairs going forward.

The other parameter we investigated was the length of a1/a2 complementarity, a region of the ncRNA structure where the 5’ and 3’ ends of the ncRNA fold back upon themselves, which we hypothesize plays a role in initiating reverse transcription (**Fig 1h**). Because this region of the ncRNA is not reverse transcribed, we cannot sequence the variants in the RT-DNA population directly. Instead, we introduced a nine base barcode in an extended loop of the msd that we could sequence as a proxy for the modification (**Fig 1i**). We amplified these barcodes directly from the purified RT-DNA for sequencing (**Fig 1j**) or prepped the RT-DNA using the TdT extension method described above (**Supplementary Fig 1b**), in both cases finding a similar effect. In this case, we found that reducing the length of complementarity in this region below seven base pairs substantially impaired RT-DNA production, consistent with a critical role in reverse transcription (**Fig 1j**). However, extending the a1/a2 length resulted in increased production of RT-DNA relative to the wt length. This is the first modification to a retron ncRNA that has been shown to increase RT-DNA yield.

### RT-DNA production in eukaryotic cells

We next wondered whether the increased production by the extended a1/a2 region would be a portable modification, to eukaryotic species and to other retrons. To facilitate expression of Eco1 in eukaryotic cells, we inverted the operon from its native arrangement. In the endogenous arrangement, the ncRNA is in the 5’ UTR of the RT transcript, requiring internal ribosome entry for the RT from an RBS that is contained in or near the a2 region of the ncRNA. In eukaryotic cells, this arrangement puts the entire ncRNA between the 5’ mRNA cap and the initiation codon for the RT. This increased distance between the cap and initiation codon, as well as the ncRNA structure and out-of-frame ATG codons, is expected to negatively affect RT translation^21^. Moreover, altering the a1/a2 region in the native arrangement could have unintended effects on the RT translation. In the inverted architecture, the RT is driven by a pol II promoter directly with its initiation codon near the 5’ end of the transcript and the ncRNA in the 3’ UTR, where variations are unlikely to influence RT translation (**Fig 2a**).

**Figure 2.**
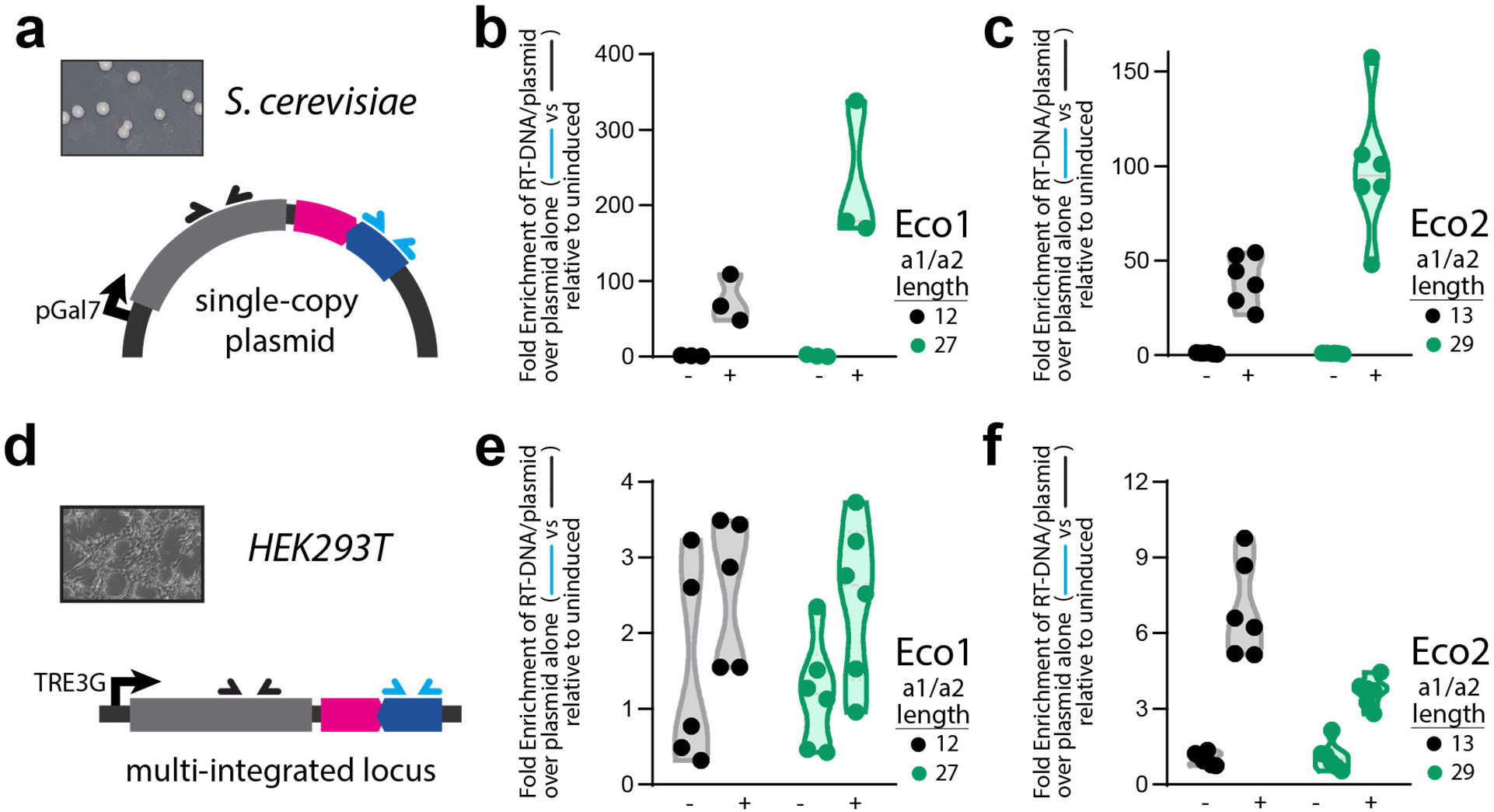
RT-DNA production in eukaryotic cells. **a.** Schematic of the retron for expression in yeast with qPCR primers indicated. **b.** Enrichment of the Eco1 RT-DNA/plasmid template over the plasmid alone by qPCR in yeast, with each construct shown relative to uninduced. Circles show each of three biological replicates, with black for the wt a1/a2 length and green for the extended a1/a2. **c.** qPCR of Eco2 in yeast, otherwise identical to b. **d.** Schematic of the retron for expression in mammalian cells with qPCR primers indicated. **e.** qPCR of Eco1 in HEK293T cells, otherwise identical to b. **f.** qPCR of Eco2 in HEK293T cells, otherwise identical to b. All statistics in Supplementary Table 1.

We first tested this arrangement for Eco1 in yeast, *S. cerevisiae.* We detected RT-DNA production using a qPCR, similar to the *E. coli* assay, comparing primers that could use the plasmid or RT-DNA as a template to primers that could only amplify the plasmid. In yeast, we found that extending the Eco1 a1/a2 region from 12 to 27 base pairs resulted in more abundant RT-DNA production (**Fig 2b, Supplementary Fig 2a**). We also extended this analysis to another retron, Eco2^19^. We found a similar effect, with the wt producing detectable RT-DNA, but a version extending the a1/a2 region from 13 to 29 base pairs producing significantly more RT-DNA (**Fig 2c, Supplementary Fig 2a**). In each case, we compared induced to uninduced cells, which likely under-reports the total RT-DNA abundance if there is any transcriptional ‘leak’ from the plasmid in the absence of inducers. Indeed, there is detectable RT-DNA in the uninduced condition relative to a control with an inactive RT, indicating some transcriptional ‘leak’ (**Supplementary Fig 2b**).

We next moved from yeast to cultured human cells, HEK293T. Using a similar gene architecture to yeast, but with a genome-integrating cassette (**Fig 2d**), we found that Eco1 does not produce significant RT-DNA in human cells, regardless of a1/a2 length (**Fig 2e**), from a well-controlled promoter (**Supplemental Fig 2c**). In contrast, Eco2 produces detectable RT-DNA, with both a wt and extended a1/a2 region (**Fig 2f**). In human cells, however, the introduction of an extended a1/a2 region diminishes, rather than enhances, production of RT-DNA. It is possible that negative transcriptional effects of the extended a1/a2 region in the human genome negate gains made in reverse transcription. Nevertheless, this is the first demonstration of RT-DNA by a retron in human cells.

### Improvements extend to applications in genome editing

In prokaryotes, retron-derived RT-DNA can be used as a template for recombineering^6, 9^. The retron ncRNA is modified to include a long loop in the msd that contains homology to a genomic locus along with one or more nucleotide modifications (**Fig 3a**). When RT-DNA from this modified ncRNA is produced along with a single stranded annealing protein (SSAP; e.g. lambda Redβ), the RT-DNA is incorporated into the lagging strand during genome replication, editing the genome of half the cell progeny in the process. This process is typically carried out in modified bacterial strains with numerous nucleases and repair proteins knocked out because editing occurs at a low rate in wt cells^9^. Therefore, we asked whether increasing RT-DNA abundance using retrons with extended a1/a2 regions could increase the rate of editing in relatively unmodified strains.

**Figure 3.**
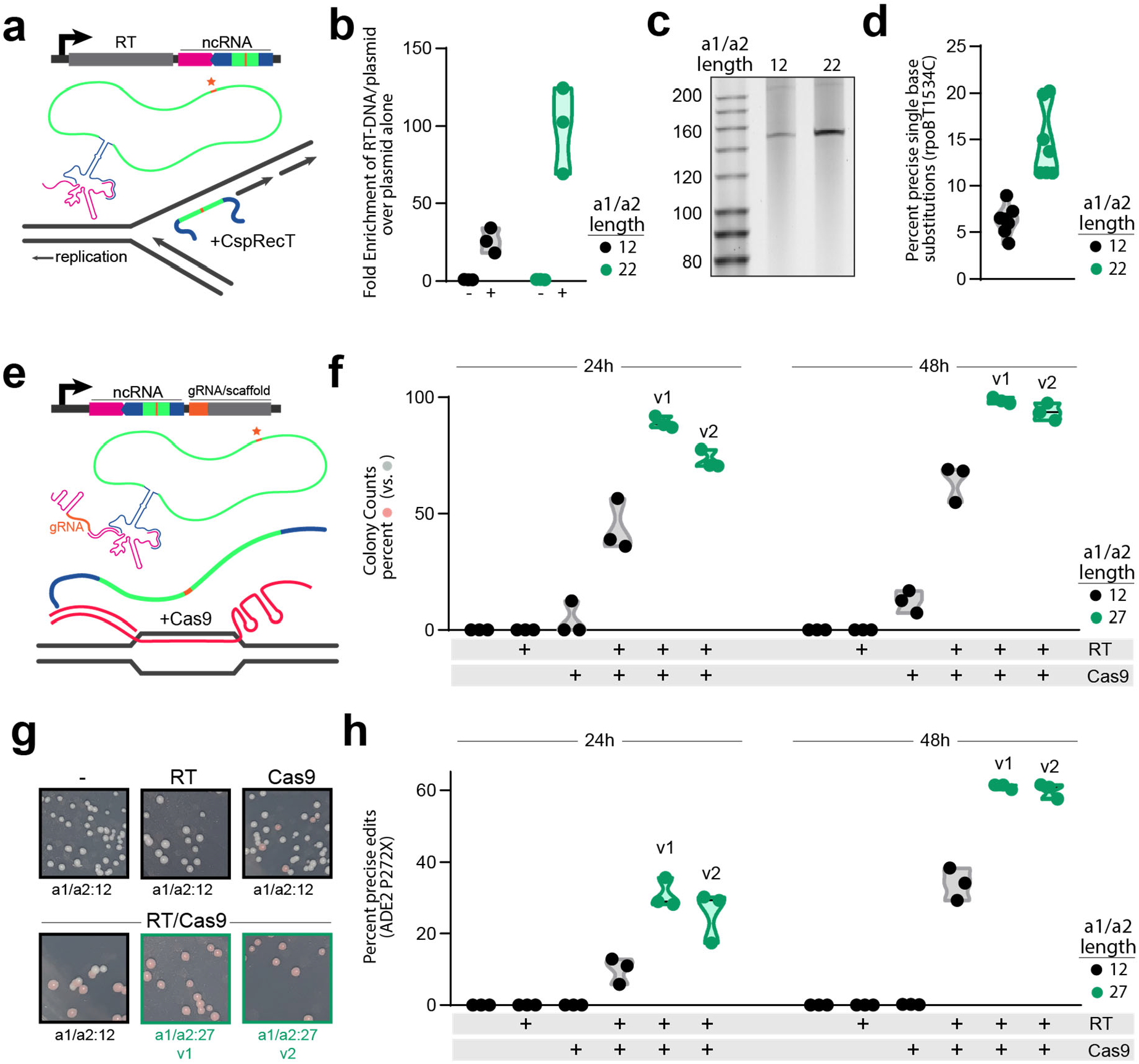
Improvements extend to applications in genome editing. **a.** Schematic of an RT-DNA template for recombineering. **b.** Enrichment of the Eco1-based recombineering RT-DNA/plasmid template over the plasmid alone by qPCR in *E. coli,* with each construct shown relative to uninduced. Circles show each of three biological replicates, with black for the wt a1/a2 length and green for the extended a1/a2. **c.** PAGE gel showing the purified RT-DNA for the wt (12) and extended (22) recombineering constructs. **d.** Percent of cells precisely edited, quantified by multiplexed sequencing, for the wt (black) and extended (green) recombineering constructs. **e.** Schematic of an RT-DNA/gRNA hybrid for genome editing in yeast. **f.** Percent of colonies edited based on phenotype (pink colonies) at 24 and 48 hours. Circles show each of three biological replicates, with black for the wt a1/a2 length and green for the extended a1/a2 (two extended versions, v1 and v2). Induction conditions are shown below the graph for the RT and Cas9. **g.** Example images from each condition plotted in f, at 24h. Induction conditions above each image. **h.** Quantification of precise editing of the ADE2 locus in yeast by Illumina sequencing, plotted as in f. All statistics in Supplementary Table 1.

We produced RT-DNA to edit a single nucleotide in the rpoB gene. We designed the retron using the same flexible architecture that we used for both the yeast and mammalian expression, with the ncRNA in the 3’ UTR of the RT. We used a 12 base stem for the msd, which retains near-wt RT-DNA production. We constructed two versions of the editing retron, one with an endogenous 12-base a1/a2 region and another with an extended 22-base a1/a2 length. Using qPCR and PAGE analysis, we confirmed that the extended a1/a2 version produced more abundant RT-DNA (**Fig 3b, c**). Finally, we expressed each version of the ncRNA along with CspRecT, a high-efficiency SSAP^22^, and mutL E32K, a dominant-negative mutL that eliminates mismatch repair at sites of single-base mismatch^23, 24^ in BL21-AI cells that were unmodified other than removal of the endogenous retron operon. Both ncRNAs resulted in appreciable editing after a single 16h overnight expression, but the extended version was significantly more effective (**Fig 3d**). This shows that the abundance of the RT-DNA template for recombineering is a limiting factor for editing, and modified ncRNA can be used to introduce edits at a higher rate.

Retron-derived RT-DNA can also be used to edit eukaryotic cells^8^. Specifically, in yeast, the ncRNA is modified to contain homology to a genomic locus and to add one or more nucleotide modifications in the loop of the msd, similar to the prokaryotic template. However, in this version, the ncRNA is on a transcript that also includes a CRISPR gRNA and scaffold. When these components are expressed along with RT and Cas9, the genomic site is cut and repaired precisely using the RT-DNA as a template (**Fig 3e**). We tested our modified ncRNAs, using an architecture that was otherwise unchanged from a previously described version^8^. The ncRNA/gRNA transcript was expressed using an inducible promoter from a plasmid, flanked by ribozyme sites. Along with the plasmid-encoded ncRNA/gRNA, we expressed either Eco1 RT, Cas9, both the RT and Cas9 or nothing at all from inducible cassettes integrated into the genome. The goal was to edit the *ADE2* locus, resulting in both a two-nucleotide edit and a cellular phenotype, pink colonies.

Using the ncRNA with a 12-base a1/a2 length, we found that the expression of both the RT and Cas9 were necessary for editing based on pink colony counts, with only a small amount of background editing when we expressed Cas9 alone (**Fig 2f,g**). This is consistent with the reverse transcription of the ncRNA being required, rather than having the edit arise from the plasmid as donor. In this case, we made two versions of the a1/a2 extended forms, which both had a length of 27 base pairs, but differed in the sequence of the a1/a2 region. We found that both versions outperformed the standard 12 base form (**Fig 2f,g**). Similar to the prokaryotic version, this indicates that the production of RT-DNA is limiting for the editing and that extended a1/a2 length is a generalizable modification that enhances retron-based genome engineering. We confirmed these phenotypic results by sequencing the *ADE2* locus from batch cultures of cells (**Fig 2h**). Precise modifications of the site, resulting from edits that use the RT-DNA as a template, follow the same pattern as the phenotypic results, showing editing that depends on both the Cas9 nuclease and RT, and is increased by extension of the a1/a2 region.

Rates of precise editing determined by sequencing from batch cultures were lower overall than editing rates estimated from counting colonies. This is likely due to additional editing that continues to occur on the plate before counting, and the requirement to score a colony as pink even if it is only partially pink. Another source could be imprecise edits to the site that result in the pink phenotype. We observed some *ADE2* loci that did not match the wt or precisely edited sequence. These occurred at a low rate (~1-3%) in all conditions, which was slightly elevated by Cas9 expression, but unaffected by RT expression/RT-DNA production (**Supplementary Fig 3a**). This, as well as the pattern of insertions, deletions, transitions, and transversions is consistent with a combination of sequencing errors and Cas9-produced in/dels (**Supplementary Fig 3b,c**).

## DISCUSSION

The bacterial retron is a molecular component that can be exploited to produce designed DNA sequences *in vivo.* Our results yield a generalizable framework for retron RT-DNA production. Specifically, we show that a minimal stem length must be maintained in the msd to yield abundant RT-DNA. We also show that there is a minimum length for the a1/a2 complementary region. Perhaps most importantly, we demonstrate that the a1/a2 region can be extended beyond its wt length to produce more abundant RT-DNA, and that increasing template abundance in both bacteria and yeast increases editing efficiency.

Importantly, these modifications are portable, both across retrons and across species. The extended a1/a2 region produces more RT-DNA using Eco1 in bacteria and both Eco1 and Eco2 in yeast. Oddly, the extended a1/a2 region did not increase RT-DNA production in cultured human cells. We speculate that this could be due to a negative transcriptional effect of the highly structured sequence, and further work will be necessary to optimize RT-DNA production in human cells. Nonetheless, we provide the first demonstration of retron-produced RT-DNA in human cells.

Retrons have been used to produce DNA templates for genome engineering^6, 8, 9^, driven by the rationale that an internally produced template eliminates problems of template availability. However, there have been no investigations of whether the RT-DNA templates are abundant enough to saturate the editing, or if even more template would lead to a yet higher rate of editing. These results establish that editing template abundance is limiting for genome editing in both bacteria and yeast, because extension of the a1/a2 region, which increases the abundance of the RT-DNA, also increases editing efficiency.

Additionally, the inverted arrangement of the retron operon, with the ncRNA in the 3’ UTR of the RT transcript, was found to produce RT-DNA in bacteria, yeast, and mammalian cells. This is the first time that a single, unifying retron architecture has been shown to be compatible with all of these host systems, simplifying comparison across kingdoms. Finally, the qPCR-based analysis of RT-DNA abundance presented here is both simpler and more quantitative than previous assays. In summary, this work represents an important advance in *in vivo* DNA synthesis that will enhance genome engineering and other biotechnological applications.

## METHODS

### Constructs and Strains

For bacterial expression, a plasmid encoding the Eco1 ncRNA and RT in that order from a T7 promoter (pSLS.436) was constructed by amplifying the retron elements from the BL21-AI genome and using Gibson assembly for integration into a backbone based on pRSFDUET1. The Eco1 RT was cloned separately into the erythromycin-inducible vector pJKR-O-mphR^25^ to generate pSLS.402. Eco1 ncRNA variants were cloned behind a T7/lac promoter in a vector based on pRSFDUET-1 with BsaI sites removed to facilitate golden gate cloning (pSLS.601), described further below. Eco1 RTs along with recombineering ncRNAs driven by T7/lac promoters (pSLS.491 and pSLS.492) were synthesized by Twist in pET-21(+).

Bacterial experiments were carried out in BL21-AI cells, or a derivative of BL21-AI cells. These cells harbor a T7 polymerase driven by a ParaB, arabinose-inducible, promoter. A knockout strain for the Eco1 operon (bSLS.114) was constructed from BL21-AI cells, using a strategy based on Datsenko and Wanner (2000)^26^ to replace the retron operon with an FRT-flanked chloramphenicol resistance cassette. The replacement cassette was amplified from pKD3, adding homology arms to the Eco1 locus. This amplicon was electroporated into BL21-AI cells expressing lambda Red genes from pKD46 and clones were isolated by selection on 40% strength chloramphenicol plates. After genotyping to confirm locus-specific insertion, the chloramphenicol cassette was excised by transient expression of FLP recombinase to leave only a FRT scar.

For yeast expression, three sets of plasmids were generated. The first set of plasmids, designed to compare the genome editing efficiency of the wt vs. extended a1/a2 length Eco1 ncRNAs, were based off of pZS.157^8^, a HIS3 yeast integrating plasmid for Galactose inducible Eco1RT and Cas9 expression (Gal1-10 promoter). Variants of pZS.157 were generated by PCR, and express either an empty cassette (pSCL.004), only Cas9 (pSCL.005), only the Eco1RT (pSCL.006), or both (pZS.157). The second set of plasmids built for the genome editing experiments are based off of pZS.165^8^, a URA3+ centromere plasmid for Galactose (Gal7) inducible expression of a modified Eco1 retron ncRNA, which consists of an Eco1 msr -ADE2- targetting gRNA chimera, flanked by HH-HDV ribozymes. An initial variant of pZS.165 was generated by cloning an IDT-synthesized gBlock consisting of an Eco1 ncRNA (a1/a2 length: 12bp), which, when reverse transcribed, encodes a 200bp Ade2 repair template to introduce a stop codon (P272X) into the ADE2 gene (pSCL.002). Two additional plasmids were generated to extend the a1/a2 region of the Eco1 ncRNA to 27bp, with variations in the a1/a2 sequence (pSCL.039 and pSCL.040). The last set of plasmids, designed to compare the levels of RT-DNA production by the different retron systems, were derived from pSCL.002. IDT-synthesized gBlocks encoding a mammalian codon-optimized Eco1RT and ncRNA (wt), a dead Eco1RT and ncRNA (wt), and a human codon-optimized Eco2RT and ncRNA (wt), were cloned into pSCL.002 by Gibson Assembly, generating pSCL.027, pSCL.031 and pSCL.017, respectively. pSCL.027 was used to generate pSCL.028 by PCR, which carries a mammalian codon optimized Eco1RT and ncRNA (extended a1/a2: 27bp). Similarly, pSC.0L17 was used to generate pSCL.034 by PCR, which carries a mammalian codon optimized Eco2RT and ncRNA (extended a1/a2: 29bp).

All yeast strains were created by LiAc/SS carrier DNA/PEG transformation^27^ of BY4742^28^. Strains for evaluating the genome editing efficiency of various retron ncRNAs were created by BY4742 integration of plasmids pZS.157, pSCL.004, pSCL.005 or pSCL.006 using KpnI-linearized plasmids for homologous recombination into the HIS3 locus. Transformants were isolated on SC-HIS plates. To evaluate effect of the length of the Eco1 ncRNA a1/a2 region on genome editing efficiency, these parental strains were transformed with episomal plasmids carrying the different retron ncRNA cassettes (pSCL.002, pSCL.039, or pSCL.040), and double transformants were isolated on SC-HIS-URA plates. The result was a set of control strains which were lacking one or both components of the genome editing machinery (i.e. Eco1RT, Cas9), and 3 strains which had all components necessary for retron-mediated genome editing but differed in the length of the Eco1 ncRNA a1/a2 region (12bp vs. 27bp).

Strains designed to compare the levels of RT-DNA production by the different retron constructs were created by transformation of plasmids pSCL.027, pSCL.037 and pSCL.028 for Eco1 (wt, wt dead RT, and extended a1/a2, respectively) into BY4742; pSCL.017 and pSCL.031 for Eco2 (wt, and extended a1/a2, respectively) into BY4742. Transformants were isolated by plating on SC-URA agar plates. Expression of proteins and ncRNAs from all yeast strains was performed in liquid SC-Ura 2% Galactose media for 24h, unless specified.

For mammalian expression, synthesized gBlocks encoding human codon optimized Eco1 and Eco2 were cloned into a PiggyBac-integrating plasmid for doxycycline-inducible human protein expression (TetOn-3G promoter). Eco1 variants are wt retron-Eco1 RT and ncRNA (pKDC.018, with a1/a2 length: 12bp), extended a1/a2 length ncRNA (pKDC.019, with a1/a2 length: 27 bp), and a dead -Eco1 RT control (pKDC.020, with a1/a2 length: 27 bp). Eco2 variants were wt retron-Eco2 RT and ncRNA (pKDC.015, with a1/a2 length: 13bp), extended a1/a2 length ncRNA (pKDC.031, with a1/a2 length: 29 bp)

Stable mammalian cell lines were created using the Lipofectamine 3000 transfection protocol (Invitrogen) and a PiggyBac transposase system. T25s of 50-70% confluent HEK293T cells were transfected using 8.3 ug of retron expression plasmids (pKDC.015, pKDC.018, pKDC.019, pKDC.020, or pKDC.031) and 4.2 ug PiggyBac transposase plasmid (pCMV-hyPBase). Stable cell lines were selected with puromycin.

Plasmids and strains are listed in Supplementary Tables 2 and 3. Primers used to generate and verify strains are listed in Supplementary Table 4. All plasmids will be made available on Addgene at the time of peer-reviewed publication.

### qPCR

qPCR analysis of RT-DNA was carried out by comparing amplification from samples using two sets of primers. One set could only use the plasmid as a template because they bound outside the msd region (outside) and the other set could use either the plasmid or RT-DNA as a template because they bound inside the msd region (inside). Results were analyzed by first taking the difference in cycle threshold (CT) between the inside and outside primer sets for each biological replicate. Next, each biological replicate ΔCT was subtracted from the average ΔCT of the control condition (e.g. uninduced). Fold change was calculated as 2^-ΔΔCT^ for each biological replicate. This fold change represents the difference in abundance of the inside versus outside template, where the presence of RT-DNA leads to fold change values >1.

For the initial analysis of Eco1 RT-DNA when overexpressed in *E. coli,* the qPCR analysis used just three primers, two of which bound inside the msd and one which bound outside. The inside PCR was generated using both inside primers, while the outside PCR used one inside and one outside primer. For all other experiments, four primers were used. Two bound inside the msd and two bound outside the msd in the RT. qPCR primers are all listed in Supplementary Table 4.

For bacterial experiments, constructs were expressed in liquid culture, shaking at 37C for 6-16 hours after which a volume of 25ul of culture was harvested, mixed with 25ul H2O, and incubated at 95C for 5 minutes. A volume of 0.3ul of this boiled culture was used as a template in 30ul reactions using a KAPA SYBR FAST qPCR mix.

For yeast experiments, single colonies were inoculated into SC-URA 2% Glucose and grown shaking overnight at 30C. To express the constructs, the overnight cultures were spun down, washed and resuspended in 1mL of water and passaged at a 1:30 dilution into SC-URA 2% Galactose, grown shaking for 24h at 30C. 250ul aliquots of the uninduced and induced cultures were collected for qPCR analysis. For qPCR sample preparation, the aliquots were spun down, resuspended in 50ul of water, and incubated at 100C for 15 minutes. The samples were then briefly spun down, placed on ice to cool, and 50ul of the supernatant was treated with Proteinase K by combining it with 29ul of water, 9ul of CutSmart buffer and 2ul of Proteinase K (NEB), followed by incubation at 56C for 30 minutes. The Proteinase K was inactivated by incubation at 95C for 10 minutes, followed by a 1.5-minute centrifugation at maximum speed (~21,000 g). The supernatant was collected and used as a template for qPCR reactions, consisting of 2.5ul of template in 10ul KAPA SYBR FAST qPCR reactions.

For mammalian experiments, retron expression in stable HEK293T cell lines was induced using 1 ug/mL doxycycline for 24h at 37C in 6-well plates. 1ml aliquots of induced and uninduced cell lines were collected for qPCR analysis. qPCR sample preparation and reaction mix followed the yeast experimental protocol.

### RT-DNA purification and PAGE analysis

To analyze RT-DNA on a PAGE gel after expression in *E. coli,* 2ml of culture were pelleted and nucleotides were prepared using a Qiagen mini prep protocol, substituting Epoch mini spin columns and buffers MX2 and MX3 for Qiagen components. Purified DNA was then treated with additional RNaseA/T1 mix (NEB) for 30 minutes at 37C and then single stranded DNA was isolated from the prep using an ssDNA/RNA Clean & Concentrator kit from Zymo Research. The purified RT-DNA was then analyzed on 10% Novex TBE-Urea Gels (Invitrogen), with a 1X TBE running buffer that was heated >80C before loading. Gels were stained with Sybr Gold (Thermo Fisher) and imaged on a Gel Doc imager (Bio-Rad).

To analyze RT-DNA on a PAGE gel after expression in *S. cerevisiae,* 5ml of overnight culture in SC-URA 2% Galactose was pelleted and RT-DNA was isolated by RNAse A/T1 treatment of the aqueous (RNA) phase after TRIzol extraction (Invitrogen), following the manufacturer’s recommendations with few modifications, as noted here. Cell pellets were resuspended in 500ul of RNA lysis buffer (100 mM EDTA pH8, 50 mM Tris-HCl pH8, 2% SDS) and incubated for 20 minutes at 85C, prior to the addition of the TRIzol reagent. The aqueous phase was chloroform-extracted twice. Following isopropanol precipitation, the RNA + RT-DNA pellet was resuspended in 265ul of TE and treated with 5ul of RNAse A/T1 + 30ul NEB2 buffer. The mixture was incubated for 25 minutes at 37C, after which the RT-DNA was re-precipitated by addition of equal volumes of isopropanol. The resulting RT-DNA was analyzed on Novex 10% TBE-Urea gels as described above.

### Variant library cloning

Eco1 ncRNA variant parts were synthesized by Agilent. Variant parts were flanked by BsaI type IIS cut sites and specific primers that allowed amplification of the sublibraries from a larger synthesis run. Random nucleotides were appended to the 3’ end of synthesized parts so that all sequences were the same length (150 bases). The vector to accept these parts (pSLS.601) was amplified with primers that also added BsaI sites, so that the ncRNA variant amplicons and amplified vector backbone could be combined into a golden gate reaction using BsaI-HFv2 and T4 ligase to generate a pool of variant plasmids at high efficiency when electroporated into a cloning strain. Variant libraries were miniprepped from the cloning strain and electroporated into the expression strain. Primers for library construction are listed in Supplementary Table 4. Variant parts are listed in Supplementary Table 5.

### Variant library expression and analysis

Eco1 ncRNA variant libraries were grown overnight and then diluted 1:500 for expression. A sample of the culture pre-expression was taken to quantify the variant plasmid library, mixed 1:1 with H2O and incubated at 95C for 5 minutes and then frozen at −20C. Constructs were expressed (arabinose and IPTG for the ncRNA, erythromycin for the RT) as the cells grew shaking at 37C for 5 hours, after which time two samples were collected. One was collected to quantify the variant plasmid library. That sample was mixed 1:1 with H2O and incubated at 95C for 5 minutes and then frozen at −20C, identically to the pre-expression sample. The other sample was collected to sequence the RT-DNA. That sample was prepared as described above for RT-DNA purification.

The two variant plasmid library samples (boiled cultures) taken before and after expression were amplified by PCR using primers flanking the ncRNA region that also contained adapters for Illumina sequencing preparation. The purified RT-DNA was prepared for sequencing by first treating with DBR1 (OriGene) to remove the branched RNA, then extending the 3’ end with a single nucleotide, dCTP, in a reaction with terminal deoxynucleotidyl transferase (TdT). This reaction was carried out in the absence of cobalt for 120 seconds at room temperature with the aim of adding only 5-10 cytosines before inactivating the TdT at 70C. A second complementary strand was then created from that extended product using Klenow Fragment (3’→5’ exo-) with a primer containing an Illumina adapter sequence, six guanines, and a non-guanine (H) anchor. Finally, Illumina adapters were ligated on at the 3’ end of the complementary strand using T4 ligase. In one variation, the loop of the RT-DNA for the a1/a2 library was amplified using Illumina adapter-containing primers in the RT-DNA, but outside the variable region from the purified RT-DNA directly. All products were indexed and sequenced on an Illumina MiSeq. Primers used for sequencing are listed in Supplementary Table 4.

Python software was custom written to extract variant counts from each plasmid and RT-DNA sample. In each case, these counts were then converted to a percentage of each library, or relative abundance (e.g. raw count for a variant over total counts for all variants). The relative abundance of a given variant in the RT-DNA sample was then divided by the relative abundance of that same variant in the plasmid library, using the average of the pre- and post-induction values, to control for differences in the abundance of each variant plasmid in the expression strain. Finally, these corrected abundance values were normalized to the average corrected abundance of the wt variant (set to 100%).

### Recombineering expression and analysis

In experiments using the retron ncRNA to edit bacterial genomes, the retron cassette was co-expressed with CspRecT and mutL E32K from the plasmid pORTMAGE-Ec1^22^ for 16 hours, shaking at 37C. After expression, a volume of 25ul of culture was harvested, mixed with 25ul H2O, and incubated at 95C for 5 minutes. A volume of 0.3ul of this boiled culture was used as a template in 30ul reactions with primers flanking the edit site in rpoB, which additionally contained adapters for Illumina sequencing preparation. These amplicons were indexed and sequenced on an Illumina MiSeq instrument and processed with custom Python software to quantify the percent of edited bases at position 1547 versus wt.

### Yeast editing expression and analysis

For yeast genome editing experiments, single colonies from strains containing variants of the Eco1 ncRNA-gRNA cassette (wt or extended a1/a2 length) and editing machinery (-/+ Cas9, -/+ Eco1RT) were grown in SC-HIS-URA 2% Raffinose for 24 hours, shaking at 30C. Cultures were passaged twice into SC-URA 2% Galactose (1:30 dilutions) for 24 hours, for a total of 48 hours of editing. At each timepoint (after 24h Raffinose, 24h Galactose, 48h Galactose), an aliquot of the cultures was harvested, diluted and plated on SC-URA low-ADE plates. Plates were incubated at 30C for 2-3 days, until visible and countable pink (ADE2 KO) and white (ADE2 WT) colonies grew. Editing efficiency was calculated in two ways. The first was by calculating the ratio of pink to total colonies on each plate for each timepoint. This counting was performed by an experimenter blind to the condition. The second was by deep sequencing of the target ADE2 locus. For this, we harvested cells from 250ul aliquots of the culture for each timepoint in PCR strips, and performed a genomic prep as follows. The pellets were resuspended in 120ul lysis buffer (see above), heated at 100C for 15 minutes and cooled on ice. 60ul of protein precipitation buffer (7.5M Ammonium Acetate) was added and the samples were gently inverted and placed at −20C for 10 minutes. The samples were then centrifuged at maximum speed for 2 minutes, and the supernatant was collected in new Eppendorf tubes. Nucleic acids were precipitated by adding equal parts ice-cold isopropanol, and incubating the samples at −20C for 10 minutes, followed by pelleting by centrifugation at maximum speed for 2 minutes. The pellets were washed twice with 200ul ice-cold 70% ethanol, and dissolved in 40ul of water. 0.5ul of the gDNA was used as template in 10ul reactions with primers flanking the edit site in ADE2, which additionally contained adapters for Illumina sequencing preparation. These amplicons were indexed and sequenced on an Illumina MiSeq instrument and processed with custom Python software to quantify the percent of P272X edits, caused by Cas9 cleavage of the target site on the ADE2 locus and repair using the Eco1 ncRNA-derived RT-DNA template.

## Data Availability

All data supporting the findings of this study are available within the article and its supplementary information, or will be made available from the authors upon request. Sequencing data associated with this study will be available in the NCBI SRA upon peer-reviewed publication.

## Code Availability

Custom code to process or analyze data from this study will be made available on github upon peer-reviewed publication.

## ACKNOWLEDGEMENTS

Work was supported by funding from the Simons Foundation Autism Research Initiative (SFARI) Bridge to Independence Award Program and the Pew Biomedical Scholars Program. S.L.S. acknowledges additional funding support from NIH/NIGMS (1DP2GM140917-01) and the L.K. Whittier Foundation. S.C.L is supported by a Berkeley Fellowship for Graduate Study. K.D.C is supported by an NSF Graduate Research Fellowship (2019247827), a UCSF Program for Breakthrough Biomedical Research New Frontiers Research Award, and a UCSF Discovery Fellowship. We thank Kathryn Claiborn for editorial assistance.

## Supplementary Figures

**Supplementary Figure 1. Related to Figure 1.**
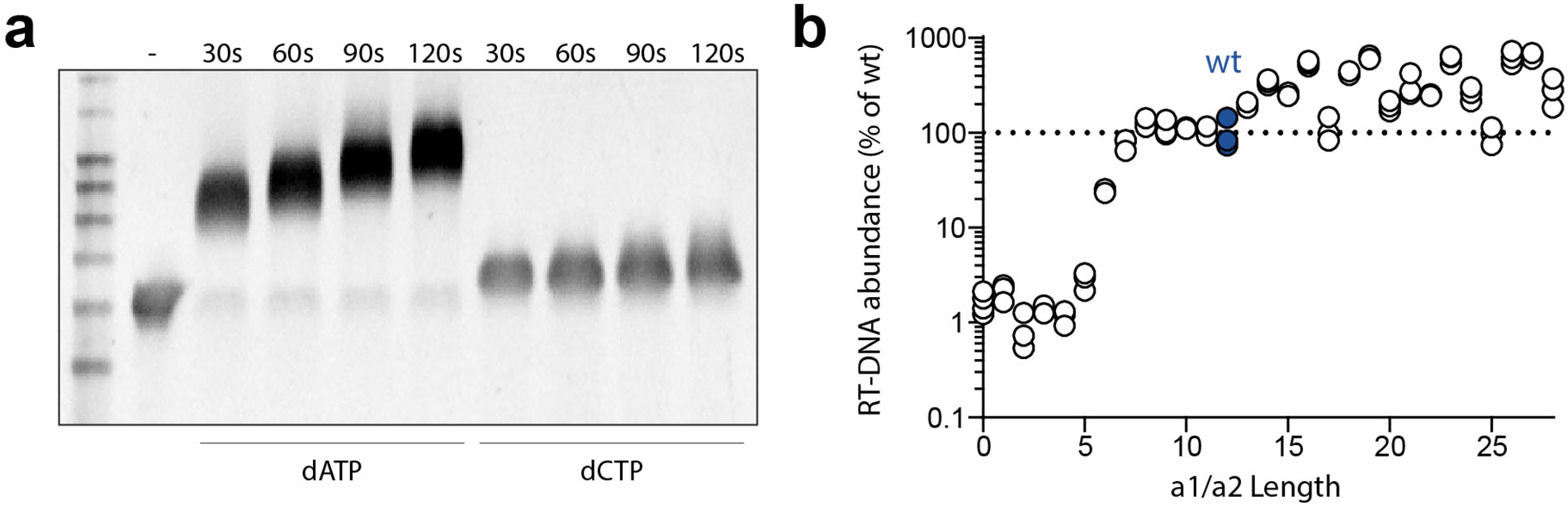
**a.** PAGE analysis showing the addition of nucleotides to the 3’ end of a single-stranded DNA, controlled by reaction time. **b.** Alternate analysis of the RT-DNA for the a1/a2 length library, using a TdT-based sequencing preparation.

**Supplementary Figure 2. Related to Figure 2.**
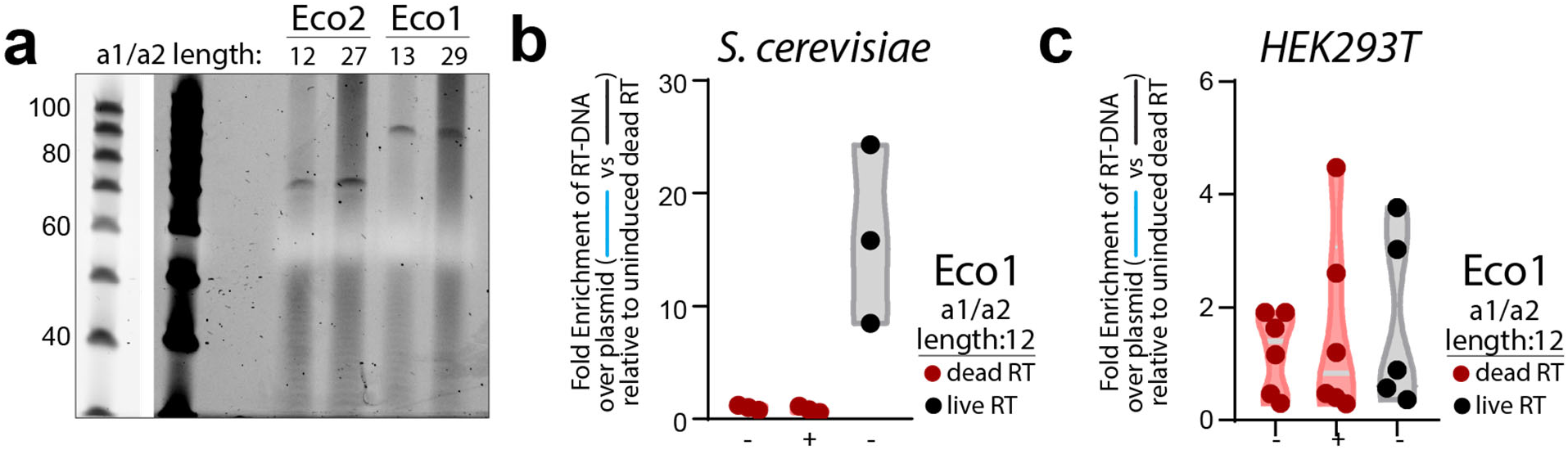
**a.** PAGE analysis of Eco1 and Eco2 RT-DNA isolated from yeast. The ladder is shown at a different exposure to the left of the gel image. **b.** Enrichment of the Eco1 RT-DNA/plasmid template when uninduced compared to a dead RT construct. Closed circles show each of three biological replicates, with red for the dead RT version and black for the live RT. **c.** Identical analysis as in b, but for Eco1 in HEK293T cells.

**Supplementary Figure 3. Related to Figure 3.**
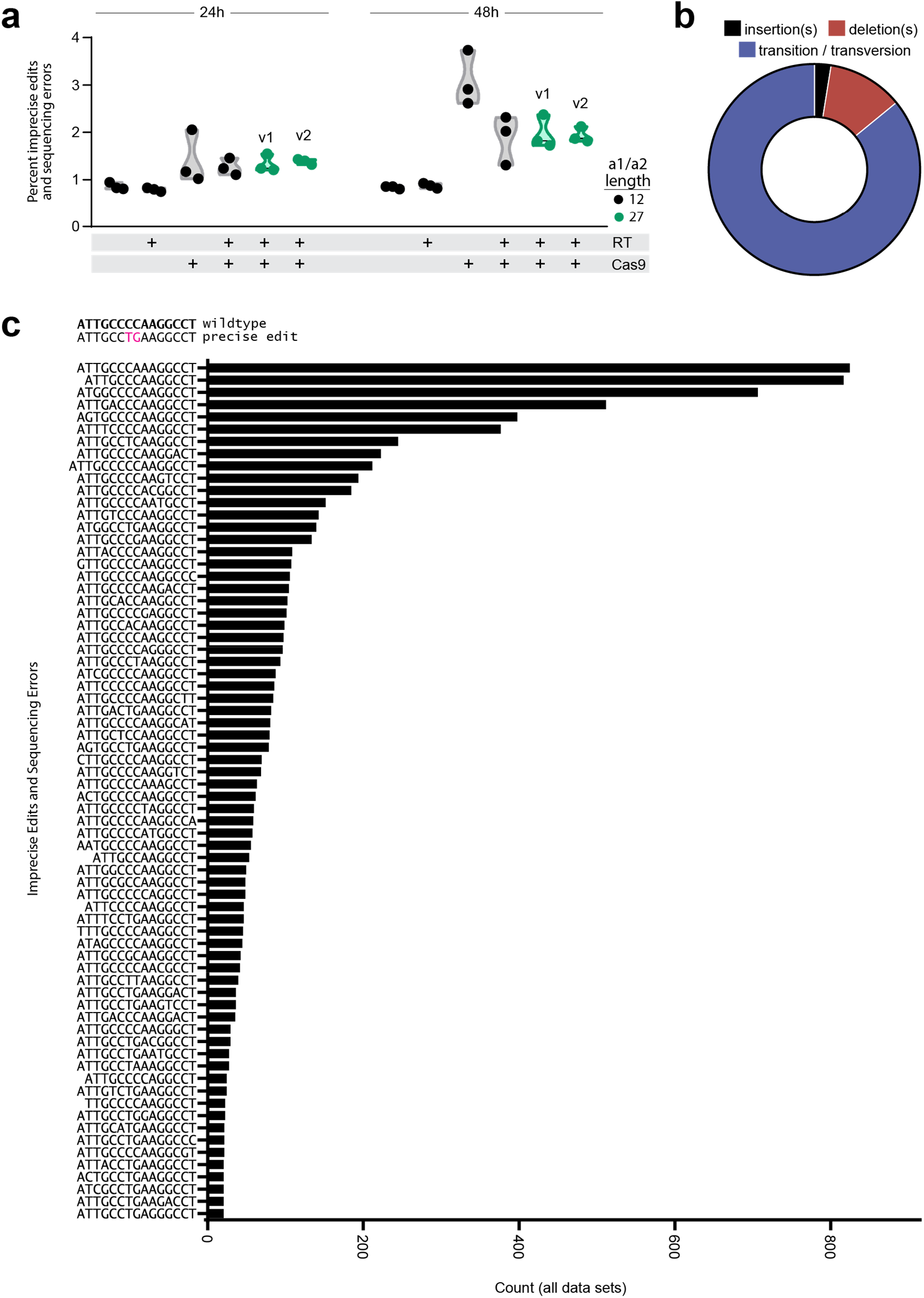
**a.** Percent of ADE2 loci with imprecise edits or sequencing errors at 24 and 48 hours. Closed circles show each of three biological replicates, with black for the wt a1/a2 length and green for the extended a1/a2 (two extended versions, v1 and v2). Induction conditions are shown below the graph for the RT and Cas9. **b.** Breakdown of the data in a. by type of edit/error. **c.** Imprecise edits and sequencing errors found in all data sets, ranked by frequency. Above the graph are the wt ADE2 locus and intended precise edit. On the Y axis are the imprecise edits and sequencing errors found. X axis represents count of each sequence in all data sets.

## References

1 Luo, D. & Saltzman, W. M. Synthetic DNA delivery systems. Nat Biotechnol 18, 33–37, doi:10.1038/71889 (2000).

2 Lin, S., Staahl, B. T., Alla, R. K. & Doudna, J. A. Enhanced homology-directed human genome engineering by controlled timing of CRISPR/Cas9 delivery. eLife 3, e04766, doi:10.7554/eLife.04766 (2014).

3 Paquet, D. et al. Efficient introduction of specific homozygous and heterozygous mutations using CRISPR/Cas9. Nature 533, 125–129, doi:10.1038/nature17664 (2016).

4 Kosicki, M., Tomberg, K. & Bradley, A. Repair of double-strand breaks induced by CRISPR-Cas9 leads to large deletions and complex rearrangements. Nat Biotechnol 36, 765–771, doi:10.1038/nbt.4192 (2018).

5 Devkota, S. The road less traveled: strategies to enhance the frequency of homology-directed repair (HDR) for increased efficiency of CRISPR/Cas-mediated transgenesis. BMB reports 51, 437–443 (2018).

6 Farzadfard, F. & Lu, T. K. Synthetic biology. Genomically encoded analog memory with precise in vivo DNA writing in living cell populations. Science 346, 1256272, doi:10.1126/science.1256272 (2014).

7 Anzalone, A. V. et al. Search-and-replace genome editing without double-strand breaks or donor DNA. Nature 576, 149–157, doi:10.1038/s41586-019-1711-4 (2019).

8 Sharon, E. et al. Functional Genetic Variants Revealed by Massively Parallel Precise Genome Editing. Cell 175, 544–557.e516, doi:10.1016/j.cell.2018.08.057 (2018).

9 Schubert, M. G. et al. High throughput functional variant screens via in-vivo production of single-stranded DNA. bioRxiv, 2020.2003.2005.975441, doi:10.1101/2020.03.05.975441 (2020).

10 Millman, A. et al. Bacterial Retrons Function In Anti-Phage Defense. Cell 183, 1551–1561.e1512, doi:10.1016/j.cell.2020.09.065 (2020).

11 Bobonis, J. et al. Bacterial retrons encode tripartite toxin/antitoxin systems. bioRxiv, 2020.2006.2022.160168, doi:10.1101/2020.06.22.160168 (2020).

12 Bobonis, J. et al. Phage proteins block and trigger retron toxin/antitoxin systems. bioRxiv, 2020.2006.2022.160242, doi:10.1101/2020.06.22.160242 (2020).

13 Gao, L. et al. Diverse enzymatic activities mediate antiviral immunity in prokaryotes. Science 369, 1077–1084, doi:10.1126/science.aba0372 (2020).

14 Mirochnitchenko, O., Inouye, S. & Inouye, M. Production of single-stranded DNA in mammalian cells by means of a bacterial retron. J Biol Chem 269, 2380–2383 (1994).

15 Lampson, B. C., Inouye, M. & Inouye, S. Retrons, msDNA, and the bacterial genome. Cytogenetic and genome research 110, 491–499, doi:10.1159/000084982 (2005).

16 Simon, A. J., Ellington, A. D. & Finkelstein, I. J. Retrons and their applications in genome engineering. Nucleic acids research 47, 11007–11019, doi:10.1093/nar/gkz865 (2019).

17 Inouye, S., Hsu, M. Y., Eagle, S. & Inouye, M. Reverse transcriptase associated with the biosynthesis of the branched RNA-linked msDNA in Myxococcus xanthus. Cell 56, 709–717, doi:10.1016/0092-8674(89)90593-x (1989).

18 Lampson, B. C., Inouye, M. & Inouye, S. Reverse transcriptase with concomitant ribonuclease H activity in the cell-free synthesis of branched RNA-linked msDNA of Myxococcus xanthus. Cell 56, 701–707, doi:10.1016/0092-8674(89)90592-8 (1989).

19 Lampson, B. C. et al. Reverse transcriptase in a clinical strain of Escherichia coli: production of branched RNA-linked msDNA. Science 243, 1033–1038 (1989).

20 Lim, D. & Maas, W. K. Reverse transcriptase-dependent synthesis of a covalently linked, branched DNA-RNA compound in E. coli B. Cell 56, 891–904 (1989).

21 Chappell, S. A., Edelman, G. M. & Mauro, V. P. Ribosomal tethering and clustering as mechanisms for translation initiation. Proceedings of the National Academy of Sciences of the United States of America 103, 18077–18082, doi:10.1073/pnas.0608212103 (2006).

22 Wannier, T. M. et al. Improved bacterial recombineering by parallelized protein discovery. Proceedings of the National Academy of Sciences of the United States of America 117, 13689–13698, doi:10.1073/pnas.2001588117 (2020).

23 Aronshtam, A. & Marinus, M. G. Dominant negative mutator mutations in the mutL gene of Escherichia coli. Nucleic acids research 24, 2498–2504, doi:10.1093/nar/24.13.2498 (1996).

24 Nyerges, Á. et al. A highly precise and portable genome engineering method allows comparison of mutational effects across bacterial species. Proceedings of the National Academy of Sciences of the United States of America 113, 2502–2507, doi:10.1073/pnas.1520040113 (2016).

25 Rogers, J. K. et al. Synthetic biosensors for precise gene control and real-time monitoring of metabolites. Nucleic acids research 43, 7648–7660, doi:10.1093/nar/gkv616 (2015).

26 Datsenko, K. A. & Wanner, B. L. One-step inactivation of chromosomal genes in Escherichia coli K-12 using PCR products. Proceedings of the National Academy of Sciences of the United States of America 97, 6640–6645, doi:10.1073/pnas.120163297 (2000).

27 Gietz, R. D. & Schiestl, R. H. High-efficiency yeast transformation using the LiAc/SS carrier DNA/PEG method. Nature protocols 2, 31–34, doi:10.1038/nprot.2007.13 (2007).

28 Brachmann, C. B. et al. Designer deletion strains derived from Saccharomyces cerevisiae S288C: a useful set of strains and plasmids for PCR-mediated gene disruption and other applications. Yeast (Chichester, England) 14, 115–132, doi:10.1002/(sici)1097-0061(19980130)14:2<115::Aid-yea204>3.0.Co;2-2 (1998).

