## Supplemental Tables for "Improved architectures for flexible DNA production using retrons across kingdoms of life"

### Supplemental Table 1, Statistics

| Panel | Biological Replicates | Comparison | Test | *P* value | *P* value summary |
|---|---|---|---|---|---|
| **Figure 1** | | | | | |
| **d** | 3 | Induced versus uninduced | unpaired t test | 0.0002 | *** |
| **g** | 3 | **Effect of stem length** | **one-way ANOVA** | **<0.0001** | **** |
| | | Follow-up: 25 vs. 0 | Dunnett's multiple comparisons test (corrected) | <0.0001 | **** |
| | | Follow-up: 25 vs. 1 | Dunnett's multiple comparisons test (corrected) | <0.0001 | **** |
| | | Follow-up: 25 vs. 2 | Dunnett's multiple comparisons test (corrected) | <0.0001 | **** |
| | | Follow-up: 25 vs. 3 | Dunnett's multiple comparisons test (corrected) | <0.0001 | **** |
| | | Follow-up: 25 vs. 4 | Dunnett's multiple comparisons test (corrected) | <0.0001 | **** |
| | | Follow-up: 25 vs. 5 | Dunnett's multiple comparisons test (corrected) | <0.0001 | **** |
| | | Follow-up: 25 vs. 6 | Dunnett's multiple comparisons test (corrected) | <0.0001 | **** |
| | | Follow-up: 25 vs. 7 | Dunnett's multiple comparisons test (corrected) | <0.0001 | **** |
| | | Follow-up: 25 vs. 8 | Dunnett's multiple comparisons test (corrected) | <0.0001 | **** |
| | | Follow-up: 25 vs. 9 | Dunnett's multiple comparisons test (corrected) | <0.0001 | **** |
| | | Follow-up: 25 vs. 10 | Dunnett's multiple comparisons test (corrected) | 0.0001 | *** |
| | | Follow-up: 25 vs. 11 | Dunnett's multiple comparisons test (corrected) | <0.0001 | **** |
| | | Follow-up: 25 vs. 12 | Dunnett's multiple comparisons test (corrected) | 0.0038 | ** |
| | | Follow-up: 25 vs. 13 | Dunnett's multiple comparisons test (corrected) | 0.1738 | ns |
| | | Follow-up: 25 vs. 14 | Dunnett's multiple comparisons test (corrected) | 0.9994 | ns |
| | | Follow-up: 25 vs. 15 | Dunnett's multiple comparisons test (corrected) | 0.0017 | ** |
| | | Follow-up: 25 vs. 16 | Dunnett's multiple comparisons test (corrected) | 0.0004 | *** |
| | | Follow-up: 25 vs. 17 | Dunnett's multiple comparisons test (corrected) | 0.657 | ns |
| | | Follow-up: 25 vs. 18 | Dunnett's multiple comparisons test (corrected) | 0.9994 | ns |
| | | Follow-up: 25 vs. 19 | Dunnett's multiple comparisons test (corrected) | 0.9997 | ns |
| | | Follow-up: 25 vs. 20 | Dunnett's multiple comparisons test (corrected) | 0.9991 | ns |
| | | Follow-up: 25 vs. 21 | Dunnett's multiple comparisons test (corrected) | 0.9997 | ns |
| | | Follow-up: 25 vs. 22 | Dunnett's multiple comparisons test (corrected) | 0.8207 | ns |
| | | Follow-up: 25 vs. 23 | Dunnett's multiple comparisons test (corrected) | 0.8984 | ns |
| | | Follow-up: 25 vs. 24 | Dunnett's multiple comparisons test (corrected) | 0.6179 | ns |
| | | Follow-up: 25 vs. 26 | Dunnett's multiple comparisons test (corrected) | 0.7321 | ns |
| | | Follow-up: 25 vs. 27 | Dunnett's multiple comparisons test (corrected) | 0.011 | * |
| | | Follow-up: 25 vs. 28 | Dunnett's multiple comparisons test (corrected) | 0.0519 | ns |
| | | Follow-up: 25 vs. 29 | Dunnett's multiple comparisons test (corrected) | 0.0489 | * |
| | | Follow-up: 25 vs. 30 | Dunnett's multiple comparisons test (corrected) | 0.0062 | ** |
| | | Follow-up: 25 vs. 31 | Dunnett's multiple comparisons test (corrected) | <0.0001 | **** |
| **j** | 3 | **Effect of a1/a2 length** | **one-way ANOVA** | **<0.0001** | **** |
| | | Follow-up: 12 vs. 0 | Dunnett's multiple comparisons test (corrected) | 0.0002 | *** |
| | | Follow-up: 12 vs. 1 | Dunnett's multiple comparisons test (corrected) | 0.0016 | ** |
| | | Follow-up: 12 vs. 2 | Dunnett's multiple comparisons test (corrected) | 0.0013 | ** |
| | | Follow-up: 12 vs. 3 | Dunnett's multiple comparisons test (corrected) | 0.0014 | ** |
| | | Follow-up: 12 vs. 4 | Dunnett's multiple comparisons test (corrected) | 0.0013 | ** |
| | | Follow-up: 12 vs. 5 | Dunnett's multiple comparisons test (corrected) | 0.0016 | ** |
| | | Follow-up: 12 vs. 6 | Dunnett's multiple comparisons test (corrected) | 0.0399 | * |
| | | Follow-up: 12 vs. 7 | Dunnett's multiple comparisons test (corrected) | 0.9944 | ns |
| | | Follow-up: 12 vs. 8 | Dunnett's multiple comparisons test (corrected) | 0.8843 | ns |
| | | Follow-up: 12 vs. 9 | Dunnett's multiple comparisons test (corrected) | 0.9988 | ns |
| | | Follow-up: 12 vs. 10 | Dunnett's multiple comparisons test (corrected) | 0.9993 | ns |
| | | Follow-up: 12 vs. 11 | Dunnett's multiple comparisons test (corrected) | 0.0559 | ns |
| | | Follow-up: 12 vs. 13 | Dunnett's multiple comparisons test (corrected) | 0.0011 | ** |
| | | Follow-up: 12 vs. 14 | Dunnett's multiple comparisons test (corrected) | <0.0001 | **** |
| | | Follow-up: 12 vs. 15 | Dunnett's multiple comparisons test (corrected) | <0.0001 | **** |
| | | Follow-up: 12 vs. 16 | Dunnett's multiple comparisons test (corrected) | <0.0001 | **** |
| | | Follow-up: 12 vs. 17 | Dunnett's multiple comparisons test (corrected) | <0.0001 | **** |
| | | Follow-up: 12 vs. 18 | Dunnett's multiple comparisons test (corrected) | <0.0001 | **** |
| | | Follow-up: 12 vs. 19 | Dunnett's multiple comparisons test (corrected) | <0.0001 | **** |
| | | Follow-up: 12 vs. 20 | Dunnett's multiple comparisons test (corrected) | 0.0107 | * |
| | | Follow-up: 12 vs. 21 | Dunnett's multiple comparisons test (corrected) | <0.0001 | **** |
| | | Follow-up: 12 vs. 22 | Dunnett's multiple comparisons test (corrected) | <0.0001 | **** |
| | | Follow-up: 12 vs. 23 | Dunnett's multiple comparisons test (corrected) | <0.0001 | **** |
| | | Follow-up: 12 vs. 24 | Dunnett's multiple comparisons test (corrected) | <0.0001 | **** |
| | | Follow-up: 12 vs. 25 | Dunnett's multiple comparisons test (corrected) | <0.0001 | **** |
| | | Follow-up: 12 vs. 26 | Dunnett's multiple comparisons test (corrected) | <0.0001 | **** |
| | | Follow-up: 12 vs. 27 | Dunnett's multiple comparisons test (corrected) | <0.0001 | **** |
| | | Follow-up: 12 vs. 28 | Dunnett's multiple comparisons test (corrected) | <0.0001 | **** |

| Panel | Biological Replicates | Comparison | Test | *P* value | *P* value summary |
|---|---|---|---|---|---|
| **Figure 2** | | | | | |
| **b** | 3 | **Effect of condition** | **one-way ANOVA** | 0.0015 | ** |
| | | Follow-up: a1/a2 length 12, induced versus uninduced | Sidak's multiple comparisons test (corrected) | 0.2898 | ns |
| | | Follow-up: a1/a2 length 27, induced versus uninduced | Sidak's multiple comparisons test (corrected) | 0.0015 | ** |
| | | Follow-up: a1/a2 length 12 versus 27, induced | Sidak's multiple comparisons test (corrected) | 0.0155 | * |

### Supplemental Table 1 continued

| | | | | | |
|---|---|---|---|---|---|
| **c** | 6 | **Effect of condition** | **one-way ANOVA** | <0.0001 | **** |
| | | Follow-up: a1/a2 length 13, induced versus uninduced | Sidak's multiple comparisons test (corrected) | 0.006 | ** |
| | | Follow-up: a1/a2 length 29, induced versus uninduced | Sidak's multiple comparisons test (corrected) | <0.0001 | **** |
| | | Follow-up: a1/a2 length 13 versus 29, induced | Sidak's multiple comparisons test (corrected) | <0.0001 | **** |
| **e** | | **Effect of condition** | **one-way ANOVA** | 0.0889 | ns |
| | | Follow-up: a1/a2 length 12, induced versus uninduced | Sidak's multiple comparisons test (corrected) | 0.2897 | ns |
| | | Follow-up: a1/a2 length 27, induced versus uninduced | Sidak's multiple comparisons test (corrected) | 0.1358 | ns |
| | | Follow-up: a1/a2 length 12 versus 27, induced | Sidak's multiple comparisons test (corrected) | 0.9957 | ns |
| **f** | | **Effect of condition** | **one-way ANOVA** | <0.0001 | **** |
| | | Follow-up: a1/a2 length 13, induced versus uninduced | Sidak's multiple comparisons test (corrected) | <0.0001 | **** |
| | | Follow-up: a1/a2 length 29, induced versus uninduced | Sidak's multiple comparisons test (corrected) | 0.0012 | ** |
| | | Follow-up: a1/a2 length 13 versus 29, induced | Sidak's multiple comparisons test (corrected) | <0.0001 | **** |
| | | | **Figure 3** | | |
| **Panel** | **Biological Replicates** | **Comparison** | **Test** | ***P* value** | ***P* value summary** |
| b | 3 | **Effect of condition** | **one-way ANOVA** | <0.0001 | **** |
| | | Follow-up: a1/a2 length 12, induced versus uninduced | Sidak's multiple comparisons test (corrected) | 0.1953 | ns |
| | | Follow-up: a1/a2 length 22, induced versus uninduced | Sidak's multiple comparisons test (corrected) | 0.0001 | *** |
| | | Follow-up: a1/a2 length 12 versus 22, induced | Sidak's multiple comparisons test (corrected) | 0.0008 | *** |
| d | 7 | a1/a2 length 12 versus 22 | unpaired t test | 0.0002 | *** |
| f | 3 | **Effect of condition (construct/induction)** | **two-way ANOVA** | **<0.0001** | **** |
| | | **Effect of time** | **two-way ANOVA** | **<0.0001** | **** |
| | | Follow-up: 24h, a1/a2 length 12, RT+ versus no expression | Sidak's multiple comparisons test (corrected) | >0.9999 | ns |
| | | Follow-up: 24h, a1/a2 length 12, Cas9+ versus no expression | Sidak's multiple comparisons test (corrected) | 0.9972 | ns |
| | | ollow-up: 24h, a1/a2 length 12, RT+/Cas9+ versus no expressic | Sidak's multiple comparisons test (corrected) | <0.0001 | **** |
| | | Follow-up: 24h, a1/a2 length 12 versus 27v1, RT+/Cas9+ | Sidak's multiple comparisons test (corrected) | <0.0001 | **** |
| | | Follow-up: 24h, a1/a2 length 12 versus 27v2, RT+/Cas9+ | Sidak's multiple comparisons test (corrected) | <0.0001 | **** |
| | | Follow-up: 48h, a1/a2 length 12, RT+ versus no expression | Sidak's multiple comparisons test (corrected) | >0.9999 | ns |
| | | Follow-up: 48h, a1/a2 length 12,Cas9+ versus no expression | Sidak's multiple comparisons test (corrected) | 0.1 | ns |
| | | ollow-up: 48h, a1/a2 length 12, RT+/Cas9+ versus no expressic | Sidak's multiple comparisons test (corrected) | <0.0001 | **** |
| | | Follow-up: 48h, a1/a2 length 12 versus 27v1, RT+/Cas9+ | Sidak's multiple comparisons test (corrected) | <0.0001 | **** |
| | | Follow-up: 48h, a1/a2 length 12 versus 27v2, RT+/Cas9+ | Sidak's multiple comparisons test (corrected) | <0.0001 | **** |
| f | 3 | **Effect of condition (construct/induction)** | **two-way ANOVA** | **<0.0001** | **** |
| | | **Effect of time** | **two-way ANOVA** | **<0.0001** | **** |
| | | Follow-up: 24h, a1/a2 length 12, RT+ versus no expression | Sidak's multiple comparisons test (corrected) | >0.9999 | ns |
| | | Follow-up: 24h, a1/a2 length 12, Cas9+ versus no expression | Sidak's multiple comparisons test (corrected) | >0.9999 | ns |
| | | ollow-up: 24h, a1/a2 length 12, RT+/Cas9+ versus no expressic | Sidak's multiple comparisons test (corrected) | 0.0077 | ** |
| | | Follow-up: 24h, a1/a2 length 12 versus 27v1, RT+/Cas9+ | Sidak's multiple comparisons test (corrected) | <0.0001 | **** |
| | | Follow-up: 24h, a1/a2 length 12 versus 27v2, RT+/Cas9+ | Sidak's multiple comparisons test (corrected) | <0.0001 | **** |
| | | Follow-up: 48h, a1/a2 length 12, RT+ versus no expression | Sidak's multiple comparisons test (corrected) | >0.9999 | ns |
| | | Follow-up: 48h, a1/a2 length 12,Cas9+ versus no expression | Sidak's multiple comparisons test (corrected) | >0.9999 | ns |
| | | ollow-up: 48h, a1/a2 length 12, RT+/Cas9+ versus no expressic | Sidak's multiple comparisons test (corrected) | <0.0001 | **** |
| | | Follow-up: 48h, a1/a2 length 12 versus 27v1, RT+/Cas9+ | Sidak's multiple comparisons test (corrected) | <0.0001 | **** |
| | | Follow-up: 48h, a1/a2 length 12 versus 27v2, RT+/Cas9+ | Sidak's multiple comparisons test (corrected) | <0.0001 | **** |

### Supplemental Table 2, Plasmids

| Name | For Expression in | Genes | Promoter | Inducer (working concentration) | Used in (panels) | Reference |
|---|---|---|---|---|---|---|
| pSLS.436 | Bacteria | Eco1: ncRNA(wt) and RT | T7 | L-arabinose (0.2% w/w) | 1d | this work |
| pSLS.402 | Bacteria | Eco1 RT | mphR | erythromycin (400uM) | 1g,j | this work |
| pSLS.601 | Bacteria | Eco1 ncRNA (variants) | T7/lac | L-arabinose (0.2% w/w) + IPTG (1mM) | 1g,j | this work |
| pSLS.491 | Bacteria | Eco1 RT and recombineering ncRNA, rpoB T1534C, a1/a2 length: 12 | T7/lac | L-arabinose (0.2% w/w) + IPTG (1mM) | 3b,c,d | this work |
| pSLS.492 | Bacteria | Eco1 RT and recombineering ncRNA, rpoB T1534C, a1/a2 length: 22 | T7/lac | L-arabinose (0.2% w/w) + IPTG (1mM) | 3b,c,d | this work |
| pORTMAGE-Ec1 | Bacteria | CspRecT and mutL E32K | Pm | m-toluic acid (1mM) | 3b,c,d | Wannier et al. (2020) |
| pKD46 | Bacteria | lambda Red genes | ParaB | L-arabinose (0.2% w/w) | - | Datsenko & Wanner (2000) |
| pKD3 | Bacteria | FRT-cat-FRT (gene disruption) | cat | - | - | Datsenko & Wanner (2000) |
| pSCL.027 | Yeast | Eco1: RT and ncRNA(wt), a1/a2 length: 12 | Gal7 | Galactose (2% w/w) | 2b | this work |
| pSCL.028 | Yeast | Eco1: RT and ncRNA(extended), a1/a2 length: 27 | Gal7 | Galactose (2% w/w) | 2b | this work |
| pSCL.037 | Yeast | Eco1: RT and ncRNA(wt), a1/a2 length: 12, dead RT | Gal7 | Galactose (2% w/w) | Supplemental 2b | this work |
| pSCL.017 | Yeast | Eco2: RT and ncRNA(wt), a1/a2 length: 13 | Gal7 | Galactose (2% w/w) | 2c | this work |
| pSCL.031 | Yeast | Eco2: RT and ncRNA(extended), a1/a2 length: 29 | Gal7 | Galactose (2% w/w) | 2c | this work |
| pSCL.004 | Yeast | Integrating inducible casette: empty | Gal1-10 | Galactose (2% w/w) | 3f,g,h | this work |
| pSCL.006 | Yeast | Integrating inducible cassette: Eco1 RT | Gal1-10 | Galactose (2% w/w) | 3f,g,h | this work |
| pSCL.005 | Yeast | Integrating inducible cassette: Cas9 | Gal1-10 | Galactose (2% w/w) | 3f,g,h | this work |
| pZS.157 | Yeast | Integrating inducible cassette: Cas9 and Eco1 RT | Gal1-10 | Galactose (2% w/w) | 3f,g,h | Sharon et al. (2018) (Addgene 114454) |
| pSCL.002 | Yeast | Eco1 editing ncRNA and gRNA, ADE2 P272X, a1/a2 length: 12 | Gal7 | Galactose (2% w/w) | 3f,g,h | reconstructed based on Sharon et al. (2018) |
| pSCL.039 | Yeast | Eco1 editing ncRNA and gRNA, ADE2 P272X, a1/a2 length: 27 v1 | Gal7 | Galactose (2% w/w) | 3f,g,h | this work |
| pSCL.040 | Yeast | Eco1 editing ncRNA and gRNA, ADE2 P272X, a1/a2 length: 27 v2 | Gal7 | Galactose (2% w/w) | 3f,g,h | this work |
| pKDC.018 | Mammalian | Eco1: RT and ncRNA(wt), a1/a2 length: 12 | TetOn-3g | Doxycycline (1 ug/ml) | 2e | this work |
| pKDC.019 | Mammalian | Eco1: RT and ncRNA(extended), a1/a2 length: 27 | TetOn-3g | Doxycycline (1 ug/ml) | 2e | this work |
| pKDC.020 | Mammalian | Eco1: RT and ncRNA(wt), a1/a2 length: 12, dead RT | TetOn-3g | Doxycycline (1 ug/ml) | Supplemental 2c | this work |
| pKDC.015 | Mammalian | Eco2: RT and ncRNA(wt), a1/a2 length: 13 | TetOn-3g | Doxycycline (1 ug/ml) | 2f | this work |
| pKDC.031 | Mammalian | Eco2: RT and ncRNA(extended), a1/a2 length: 29 | TetOn-3g | Doxycycline (1 ug/ml) | 2f | this work |

### Supplemental Table 3, Strains

| Name | Species | Parental Line | Genotype | Method |
|---|---|---|---|---|
| bSLS.114 | *E. coli* | BL21-AI | Eco1 KO | lambda Red recombinase mediated insertion of chloramphenicol resistance, marker excision by FLP |
| ySCL2 | *S. cerevisiae* | BY4742 | HIS3: pSCL.004 | LiAc mediated plasmid transformation and HR mediated insertion of empty Galactose inducible cassette (pSCL.004) into the HIS3 locus |
| ySCL3 | *S. cerevisiae* | BY4742 | HIS3: pSCL.005 | LiAc mediated plasmid transformation and HR mediated insertion of a Galactose inducible cassette for Cas9 expression (pSCL.005) into the HIS3 locus [illegible] |
| ySCL4 | *S. cerevisiae* | BY4742 | HIS3: pSCL.006 | LiAc mediated plasmid transformation and HR mediated insertion of a Galactose inducible cassette for Eco1RT expression (pSCL.006) into the HIS3 loc[illegible] |
| ySCL5 | *S. cerevisiae* | BY4742 | HIS3: pZS.157 | LiAc mediated plasmid transformation and HR mediated insertion of a Galactose inducible cassette for Cas9 and Eco1RT expression (pZS.157) into the [illegible] |
| pKDC.018 | *H. sapien* | HEK293T | Integrated pKDC.018 | Lipofectamine 3000 mediated plasmid transfection and PiggyBac mediated insertion of doxycycline inducible casette for Eco1RT and wt nCRNA |
| pKDC.019 | *H. sapien* | HEK293T | Integrated pKDC.019 | Lipofectamine 3000 mediated plasmid transfection and PiggyBac mediated insertion of doxycycline inducible casette for Eco1RT and extended a1/a2 [illegible] |
| pKDC.020 | *H. sapien* | HEK293T | Integrated pKDC.020 | Lipofectamine 3000 mediated plasmid transfection and PiggyBac mediated insertion of doxycycline inducible casette for Eco1 deadRT and extended a[illegible] |
| pKDC.015 | *H. sapien* | HEK293T | Integrated pKDC.015 | Lipofectamine 3000 mediated plasmid transfection and PiggyBac mediated insertion of doxycycline inducible casette for Eco2RT and wt nCRNA |
| pKDC.031 | *H. sapien* | HEK293T | Integrated pKDC.031 | Lipofectamine 3000 mediated plasmid transfection and PiggyBac mediated insertion of doxycycline inducible casette for Eco2RT and extended a1/a2 [illegible] |

### Supplemental Table 4, Primers

| Name | Sequence | Purpose |
|---|---|---|
| pKD3_Eco1_KO_F | GGATCTATTCATAACTTGATGTATAAAGTAGAAAAAAAAGCGGGGAGATTGTGTAGGCTGGAGCTGCTTC | Generating a knockout template for the Eco1 locus from pKD3 |
| pKD3_Eco1_KO_R | GAAGTGCTCAATTTTTCAACCTATGAGCTTTAGTTTTAAAACGAAGACCACATATGAATATCCTCCTTA | |
| Eco1_KO_genotyping_F | CATGTGCATGAAAACCACTGC | Verifying locus-specific insertion of CM cassette |
| Eco1_KO_genotyping_R | ATCATGCCGTTTGTGATGG | |
| Eco1_KO_CM_marker_loss_R | GCAATGCGATGAAAAGTGCC | With Eco1_KO_genotyping_F, verifying excision of CM cassette |
| qPCR_Eco1_wt_1 | GTCAGAAAAAACGGGTTTCCTGGTTGG | qPCR quantification of RT-DNA from Eco1 (wt) |
| qPCR_Eco1_wt_2 | TCTGAGTTACTGTCTGTTTTCCTTGTTGG | with qPCR_Eco1_wt_1, amplifies the RT-DNA and plasmid |
| qPCR_Eco1_wt_3 | CCTCTGGATGTTGTTTCGGCA | with qPCR_Eco1_wt_1, amplifies only the plasmid |
| qPCR_Eco1_Recombineering_RT_F | CAAGGATGGGTTCTACGTAACA | qPCR quantification of RT-DNA from Eco1 Recombineering ncRNA amplifies only plasmid (RT) |
| qPCR_Eco1_Recombineering_RT_R | GCTAATTTAGGTGATGATGGAGC | |
| qPCR_Eco1_Recombineering_RTDNA_F | CTGAGTTACTGTCTGTTTTCCTG | qPCR quantification of RT-DNA from Eco1 Recombineering ncRNA amplifies plasmid and RT-DNA (RT-DNA) |
| qPCR_Eco1_Recombineering_RTDNA_R | TCAGAAAAAACGGGTTTCCTGAATTC | |
| rpoB_genotyping_F | CTTTCCCTACACGACGCTCTTCCGATCTCTCTGGGCGATCTGGATACC | Genotyping of single base change in rpoB gene by Illumina sequencing |
| rpoB_genotyping_R | GGAGTTCAGACGTGTGCTCTTCCGATCTTTCGATTGGACATACGCGAC | |
| Vector_BB_for_stem_library_F | agctaGGTCTCATTCAATGCAGGATGCCGAAACAAC | Amplifies vector for insertion of stem library parts |
| Vector_BB_for_stem_library_R | agctaGGTCTCAGTAAGGGTGCGCAACTTTCATG | |
| Vector_BB_for_a1/a2_library_F | agctaGGTCTCATTCAATGCAGGATGCCGAAACAAC | Amplifies vector for insertion of a1/a2 library parts |
| Vector_BB_for_a1/a2_library_R | agctaGGTCTCAACTTTCATGAAATCCGCTGCATCAC | |
| Stem_Library_part_F | AGTGACCCGTCCCTG | Amplifies stem library variant parts |
| Stem_Library_part_R | AGTCGACCTCTGCCC | |
| a1/a2_Library_part_F | ACTGGTGCGTCGTCT | Amplifies a1/a2 library variant parts |
| a1/a2_Library_part_R | CGGGAAGTGTTCGCC | |
| Eco1_Variant_Plasmids_for_Sequencing_F | CTTTCCCTACACGACGCTCTTCCGATCTNNNNNTTATGCTAGGTGATGCAGCGGATTTCATGAAAG | Amplifies ncRNA from variant library plasmids for Illumina sequencing |
| Eco1_Variant_Plasmids_for_Sequencing_R | GGAGTTCAGACGTGTGCTCTTCCGATCTCATTTAACTATGATAAGATTCCGTATGCGCACC | |
| ssExt_pG6_anchor | GGAGTTCAGACGTGTGCTCTTCCGATCTGGGGGGH | Creates complementary strand from extended RT-DNA |
| Eco1_msdloop_for_Sequencing_F | CTTTCCCTACACGACGCTCTTCCGATCTTCAGAAAAAACGGGTTGTCGCC | Amplifies barcodes in the msd loop from purified RT-DNA for Illumina sequencing |
| Eco1_msdloop_for_Sequencing_R | GGAGTTCAGACGTGTGCTCTTCCGATCTGTTACAAGCTGTTTGTCGCCAG | |
| SCL325 | GTCGAAAAGAAGGGGCCTGA | qPCR primers to amplify the Eco1RT (yeast version) |
| SCL326 | CCGCGTACCCTTGTATTCGA | |
| SCL287 | TCCTTGTTGGAACGGAGAGC | qPCR primers to amplify the Eco1 RT-DNA (yeast version) |
| SCL288 | TAACGGGTTTCCTGGTTGGC | |
| SCL196 | ACCTTAAAGCTGCCCTCCAT | qPCR primers to amplify the Eco2RT (yeast and human version) |
| SCL197 | GGTACGCTGCCGTAATAGGA | |
| SCL34 | GAATCGCCTCCCTAAAATCC | qPCR primers to amplify the Eco2 RT-DNA (yeast and human version) |
| SCL37 | GCACACCTGCCGTATAGCTC | |
| SCL378 | CTTTCCCTACACGACGCTCTTCCGATCTTATGCGCCTGCTAGAGTTCC | Amplify the Ade2 locus for Illumina sequencing |
| SCL193 | GGAGTTCAGACGTGTGCTCTTCCGATCTGCGTTCGTTGTAATGGTGGAG | |
| SCL_gblock_001 | GAATCTGAGTTACTGTCTGTTTTCCTTGGAAATGTTCTATTTAGAAACAGGGGAATTGCTTATTAACGAAATTGCCTGA | Cloned into pZS.165 to make pSCL.002 -- ade2hdr-sgAde2 oligo: edits the ade2 locus |
| Eco1_RT_fow | ACGTTCCGCCTTAGGAATTT | qPCR primers to amplify the Eco1RT (human version) |
| Eco1_RT_rev | TTTCTCAGGCCCCTTCTTTT | |
| Eco1_RTDNA_fow | AAATAACGGGTTTCCTGGTTG | qPCR primers to amplify the Eco1 RT-DNA (human version) |
| Eco1_RTDNA_rev | TCTGTTTTCCTTGTTGGAACG | |

### Supplemental Table 5, Variant Parts

| Library | Length (stem or a1/a2) | ID | ncRNA | S |
|---|---|---|---|---|
| Eco1 Stem Length | 0 | 4235 | TGCGCACCCTTAGCGAGAGGTTTATCATTAAGGTCAACCTCTGGATGTTGTTTCGGCATCCTGCATTGAATCTGAGTTACTGTCTGTGCCTCCCGTTTTTTCTGACGTAAGGGTGCGCA | A |
| Eco1 Stem Length | 1 | 4236 | TGCGCACCCTTAGCGAGAGGTTTATCATTAAGGTCAACCTCTGGATGTTGTTTCGGCATCCTGCATTGAATCTGAGTTACTGTCTGTTGCCTACCCGTTTTTTCTGACGTAAGGGTGCGCA | A |
| Eco1 Stem Length | 2 | 4237 | TGCGCACCCTTAGCGAGAGGTTTATCATTAAGGTCAACCTCTGGATGTTGTTTCGGCATCCTGCATTGAATCTGAGTTACTGTCTGTTTGCCTAACCCGTTTTTTCTGACGTAAGGGTGCGCA | A |
| Eco1 Stem Length | 3 | 4238 | TGCGCACCCTTAGCGAGAGGTTTATCATTAAGGTCAACCTCTGGATGTTGTTTCGGCATCCTGCATTGAATCTGAGTTACTGTCTGTTTTGCCTAAACCCGTTTTTTCTGACGTAAGGGTGCGCA | A |
| Eco1 Stem Length | 4 | 4239 | TGCGCACCCTTAGCGAGAGGTTTATCATTAAGGTCAACCTCTGGATGTTGTTTCGGCATCCTGCATTGAATCTGAGTTACTGTCTGTTTTCGCCTGAAACCCGTTTTTTCTGACGTAAGGGTGCGCA | A |
| Eco1 Stem Length | 5 | 4240 | TGCGCACCCTTAGCGAGAGGTTTATCATTAAGGTCAACCTCTGGATGTTGTTTCGGCATCCTGCATTGAATCTGAGTTACTGTCTGTTTTCCGCCTGGAAACCCGTTTTTTCTGACGTAAGGGTGCGCA | A |
| Eco1 Stem Length | 6 | 4241 | TGCGCACCCTTAGCGAGAGGTTTATCATTAAGGTCAACCTCTGGATGTTGTTTCGGCATCCTGCATTGAATCTGAGTTACTGTCTGTTTTCCTGCCTAGGAAACCCGTTTTTTCTGACGTAAGGGTGCGCA | A |
| Eco1 Stem Length | 7 | 4242 | TGCGCACCCTTAGCGAGAGGTTTATCATTAAGGTCAACCTCTGGATGTTGTTTCGGCATCCTGCATTGAATCTGAGTTACTGTCTGTTTTCCTTGCCTCAGGAAACCCGTTTTTTCTGACGTAAGGGTGCGCA | A |
| Eco1 Stem Length | 8 | 4243 | TGCGCACCCTTAGCGAGAGGTTTATCATTAAGGTCAACCTCTGGATGTTGTTTCGGCATCCTGCATTGAATCTGAGTTACTGTCTGTTTTCCTTGGCCTCCAGGAAACCCGTTTTTTCTGACGTAAGGGTGCGCA | A |
| Eco1 Stem Length | 9 | 4244 | TGCGCACCCTTAGCGAGAGGTTTATCATTAAGGTCAACCTCTGGATGTTGTTTCGGCATCCTGCATTGAATCTGAGTTACTGTCTGTTTTCCTTGTGCCTACCAGGAAACCCGTTTTTTCTGACGTAAGGGTGCGCA | A |
| Eco1 Stem Length | 10 | 4245 | TGCGCACCCTTAGCGAGAGGTTTATCATTAAGGTCAACCTCTGGATGTTGTTTCGGCATCCTGCATTGAATCTGAGTTACTGTCTGTTTTCCTTGTTGCCTAACCAGGAAACCCGTTTTTTCTGACGTAAGGGTGCGCA | A |
| Eco1 Stem Length | 11 | 4246 | TGCGCACCCTTAGCGAGAGGTTTATCATTAAGGTCAACCTCTGGATGTTGTTTCGGCATCCTGCATTGAATCTGAGTTACTGTCTGTTTTCCTTGTTGGCCTCAACCAGGAAACCCGTTTTTTCTGACGTAAGGGTGCGCA | A |
| Eco1 Stem Length | 12 | 4247 | TGCGCACCCTTAGCGAGAGGTTTATCATTAAGGTCAACCTCTGGATGTTGTTTCGGCATCCTGCATTGAATCTGAGTTACTGTCTGTTTTCCTTGTTGGGCCTCCAACCAGGAAACCCGTTTTTTCTGACGTAAGGGTGCGCA | A |
| Eco1 Stem Length | 13 | 4248 | TGCGCACCCTTAGCGAGAGGTTTATCATTAAGGTCAACCTCTGGATGTTGTTTCGGCATCCTGCATTGAATCTGAGTTACTGTCTGTTTTCCTTGTTGGAGCCTGCCAACCAGGAAACCCGTTTTTTCTGACGTAAGGGTGCGCA | A |
| Eco1 Stem Length | 14 | 4249 | TGCGCACCCTTAGCGAGAGGTTTATCATTAAGGTCAACCTCTGGATGTTGTTTCGGCATCCTGCATTGAATCTGAGTTACTGTCTGTTTTCCTTGTTGGAAGCCTAGCCAACCAGGAAACCCGTTTTTTCTGACGTAAGGGTGCGCA | A |
| Eco1 Stem Length | 15 | 4250 | TGCGCACCCTTAGCGAGAGGTTTATCATTAAGGTCAACCTCTGGATGTTGTTTCGGCATCCTGCATTGAATCTGAGTTACTGTCTGTTTTCCTTGTTGGAACGCCTGAGCCAACCAGGAAACCCGTTTTTTCTGACGTAAGGGTGCGCA | A |
| Eco1 Stem Length | 16 | 4251 | TGCGCACCCTTAGCGAGAGGTTTATCATTAAGGTCAACCTCTGGATGTTGTTTCGGCATCCTGCATTGAATCTGAGTTACTGTCTGTTTTCCTTGTTGGAACGGCCTCGAGCCAACCAGGAAACCCGTTTTTTCTGACGTAAGGGTGCGCA | A |
| Eco1 Stem Length | 17 | 4252 | TGCGCACCCTTAGCGAGAGGTTTATCATTAAGGTCAACCTCTGGATGTTGTTTCGGCATCCTGCATTGAATCTGAGTTACTGTCTGTTTTCCTTGTTGGAACGGGCCTCCGAGCCAACCAGGAAACCCGTTTTTTCTGACGTAAGGGTGCGCA | A |
| Eco1 Stem Length | 18 | 4253 | TGCGCACCCTTAGCGAGAGGTTTATCATTAAGGTCAACCTCTGGATGTTGTTTCGGCATCCTGCATTGAATCTGAGTTACTGTCTGTTTTCCTTGTTGGAACGGAGCCTTCCGAGCCAACCAGGAAACCCGTTTTTTCTGACGTAAGGGTGCGCA | A |
| Eco1 Stem Length | 19 | 4254 | TGCGCACCCTTAGCGAGAGGTTTATCATTAAGGTCAACCTCTGGATGTTGTTTCGGCATCCTGCATTGAATCTGAGTTACTGTCTGTTTTCCTTGTTGGAACGGAGGCCTCTCCGAGCCAACCAGGAAACCCGTTTTTTCTGACGTAAGGGTGCGCA | A |
| Eco1 Stem Length | 20 | 4255 | TGCGCACCCTTAGCGAGAGGTTTATCATTAAGGTCAACCTCTGGATGTTGTTTCGGCATCCTGCATTGAATCTGAGTTACTGTCTGTTTTCCTTGTTGGAACGGAGAGCCTTCTCCGAGCCAACCAGGAAACCCGTTTTTTCTGACGTAAGGGTGCGCA | A |
| Eco1 Stem Length | 21 | 4256 | TGCGCACCCTTAGCGAGAGGTTTATCATTAAGGTCAACCTCTGGATGTTGTTTCGGCATCCTGCATTGAATCTGAGTTACTGTCTGTTTTCCTTGTTGGAACGGAGAGGCCTCTCTCCGAGCCAACCAGGAAACCCGTTTTTTCTGACGTAAGGGTGCGCA | A |
| Eco1 Stem Length | 22 | 4257 | TGCGCACCCTTAGCGAGAGGTTTATCATTAAGGTCAACCTCTGGATGTTGTTTCGGCATCCTGCATTGAATCTGAGTTACTGTCTGTTTTCCTTGTTGGAACGGAGAGCGCCTGCTCTCCGAGCCAACCAGGAAACCCGTTTTTTCTGACGTAAGGGTGCGCA | A |
| Eco1 Stem Length | 23 | 4258 | TGCGCACCCTTAGCGAGAGGTTTATCATTAAGGTCAACCTCTGGATGTTGTTTCGGCATCCTGCATTGAATCTGAGTTACTGTCTGTTTTCCTTGTTGGAACGGAGAGCAGCCTTGCTCTCCGAGCCAACCAGGAAACCCGTTTTTTCTGACGTAAGGGTGCGCA | A |
| Eco1 Stem Length | 24 | 4259 | TGCGCACCCTTAGCGAGAGGTTTATCATTAAGGTCAACCTCTGGATGTTGTTTCGGCATCCTGCATTGAATCTGAGTTACTGTCTGTTTTCCTTGTTGGAACGGAGAGCATGCCTATGCTCTCCGAGCCAACCAGGAAACCCGTTTTTTCTGACGTAAGGGTGCGCA | A |
| Eco1 Stem Length | 25 | 4260 | TGCGCACCCTTAGCGAGAGGTTTATCATTAAGGTCAACCTCTGGATGTTGTTTCGGCATCCTGCATTGAATCTGAGTTACTGTCTGTTTTCCTTGTTGGAACGGAGAGCATCGCCTGATGCTCTCCGAGCCAACCAGGAAACCCGTTTTTTCTGACGTAAGGGTGCGCA | A |
| Eco1 Stem Length | 26 | 4261 | TGCGCACCCTTAGCGAGAGGTTTATCATTAAGGTCAACCTCTGGATGTTGTTTCGGCATCCTGCATTGAATCTGAGTTACTGTCTGTTTTCCTTGTTGGAACGGAGAGCATCGGCCTCGATGCTCTCCGAGCCAACCAGGAAACCCGTTTTTTCTGACGTAAGGGTGCGCA | A |
| Eco1 Stem Length | 27 | 4262 | TGCGCACCCTTAGCGAGAGGTTTATCATTAAGGTCAACCTCTGGATGTTGTTTCGGCATCCTGCATTGAATCTGAGTTACTGTCTGTTTTCCTTGTTGGAACGGAGAGCATCTAGCCTTAGATGCTCTCCGAGCCAACCAGGAAACCCGTTTTTTCTGACGTAAGGGTGCGCA | A |
| Eco1 Stem Length | 28 | 4263 | TGCGCACCCTTAGCGAGAGGTTTATCATTAAGGTCAACCTCTGGATGTTGTTTCGGCATCCTGCATTGAATCTGAGTTACTGTCTGTTTTCCTTGTTGGAACGGAGAGCATCAGTGCCTACTGATGCTCTCCGAGCCAACCAGGAAACCCGTTTTTTCTGACGTAAGGGTGCGCA | A |
| Eco1 Stem Length | 29 | 4264 | TGCGCACCCTTAGCGAGAGGTTTATCATTAAGGTCAACCTCTGGATGTTGTTTCGGCATCCTGCATTGAATCTGAGTTACTGTCTGTTTTCCTTGTTGGAACGGAGAGCATCTCACGCCTGTGAGATGCTCTCCGAGCCAACCAGGAAACCCGTTTTTTCTGACGTAAGGGTGCGCA | A |
| Eco1 Stem Length | 30 | 4265 | TGCGCACCCTTAGCGAGAGGTTTATCATTAAGGTCAACCTCTGGATGTTGTTTCGGCATCCTGCATTGAATCTGAGTTACTGTCTGTTTTCCTTGTTGGAACGGAGAGCATCCGAATGCCTATTCGGATGCTCTCCGAGCCAACCAGGAAACCCGTTTTTTCTGACGTAAGGGTGCGCA | A |
| Eco1 Stem Length | 31 | 4266 | TGCGCACCCTTAGCGAGAGGTTTATCATTAAGGTCAACCTCTGGATGTTGTTTCGGCATCCTGCATTGAATCTGAGTTACTGTCTGTTTTCCTTGTTGGAACGGAGAGCATCAAAACGGCCTCGTTTTGATGCTCTCCGAGCCAACCAGGAAACCCGTTTTTTCTGACGTAAGGGTGCGCA | A |
| Eco1 a1/a2 Length | 0 | 5267 | CGAGAGGTTTATCATTAAGGTCAACCTCTGGATGTTGTTTCGGCATCCTGCATTGAATCTGAGTTACTGTCTGTTTTCCTTGTTGGAAGAACTGCTGAGCCAACCAGGAAACCCGTTTTTTCTGAC | A |
| Eco1 a1/a2 Length | 0 | 5268 | GCGAGAGGTTTATCATTAAGGTCAACCTCTGGATGTTGTTTCGGCATCCTGCATTGAATCTGAGTTACTGTCTGTTTTCCTTGTTGGAAATTAGTTTCAGCCAACCAGGAAACCCGTTTTTTCTGACG | A |
| Eco1 a1/a2 Length | 1 | 5269 | AGCGAGAGGTTTATCATTAAGGTCAACCTCTGGATGTTGTTTCGGCATCCTGCATTGAATCTGAGTTACTGTCTGTTTTCCTTGTTGGAAGATATCATCAGCCAACCAGGAAACCCGTTTTTTCTGACGT | A |
| Eco1 a1/a2 Length | 2 | 5270 | TAGCGAGAGGTTTATCATTAAGGTCAACCTCTGGATGTTGTTTCGGCATCCTGCATTGAATCTGAGTTACTGTCTGTTTTCCTTGTTGGAAACCCTAAAGAGCCAACCAGGAAACCCGTTTTTTCTGACGTA | A |
| Eco1 a1/a2 Length | 3 | 5271 | TTAGCGAGAGGTTTATCATTAAGGTCAACCTCTGGATGTTGTTTCGGCATCCTGCATTGAATCTGAGTTACTGTCTGTTTTCCTTGTTGGAAGGAAGAAAAAGCCAACCAGGAAACCCGTTTTTTCTGACGTAA | A |
| Eco1 a1/a2 Length | 4 | 5272 | CTTAGCGAGAGGTTTATCATTAAGGTCAACCTCTGGATGTTGTTTCGGCATCCTGCATTGAATCTGAGTTACTGTCTGTTTTCCTTGTTGGAAGAGGTTTCAAGCCAACCAGGAAACCCGTTTTTTCTGACGTAAG | A |
| Eco1 a1/a2 Length | 5 | 5273 | CCTTAGCGAGAGGTTTATCATTAAGGTCAACCTCTGGATGTTGTTTCGGCATCCTGCATTGAATCTGAGTTACTGTCTGTTTTCCTTGTTGGAATTTGCGCCAAGCCAACCAGGAAACCCGTTTTTTCTGACGTAAGG | A |
| Eco1 a1/a2 Length | 6 | 5274 | CCCTTAGCGAGAGGTTTATCATTAAGGTCAACCTCTGGATGTTGTTTCGGCATCCTGCATTGAATCTGAGTTACTGTCTGTTTTCCTTGTTGGAAACGAATATAAGCCAACCAGGAAACCCGTTTTTTCTGACGTAAGGG | A |
| Eco1 a1/a2 Length | 7 | 5275 | ACCCTTAGCGAGAGGTTTATCATTAAGGTCAACCTCTGGATGTTGTTTCGGCATCCTGCATTGAATCTGAGTTACTGTCTGTTTTCCTTGTTGGAACTTCGCACTAGCCAACCAGGAAACCCGTTTTTTCTGACGTAAGGGT | A |
| Eco1 a1/a2 Length | 8 | 5276 | CACCCTTAGCGAGAGGTTTATCATTAAGGTCAACCTCTGGATGTTGTTTCGGCATCCTGCATTGAATCTGAGTTACTGTCTGTTTTCCTTGTTGGAAGGAGGGCTTAGCCAACCAGGAAACCCGTTTTTTCTGACGTAAGGGTG | A |
| Eco1 a1/a2 Length | 9 | 5277 | GCACCCTTAGCGAGAGGTTTATCATTAAGGTCAACCTCTGGATGTTGTTTCGGCATCCTGCATTGAATCTGAGTTACTGTCTGTTTTCCTTGTTGGAAGAAGCAGAGAGCCAACCAGGAAACCCGTTTTTTCTGACGTAAGGGTGC | A |
| Eco1 a1/a2 Length | 10 | 5278 | CGCACCCTTAGCGAGAGGTTTATCATTAAGGTCAACCTCTGGATGTTGTTTCGGCATCCTGCATTGAATCTGAGTTACTGTCTGTTTTCCTTGTTGGAACAGGGCTGAAGCCAACCAGGAAACCCGTTTTTTCTGACGTAAGGGTGCG | A |
| Eco1 a1/a2 Length | 11 | 5279 | GCGCACCCTTAGCGAGAGGTTTATCATTAAGGTCAACCTCTGGATGTTGTTTCGGCATCCTGCATTGAATCTGAGTTACTGTCTGTTTTCCTTGTTGGAAAAAACTGTCAGCCAACCAGGAAACCCGTTTTTTCTGACGTAAGGGTGCGC | A |
| Eco1 a1/a2 Length | 12 | 5280 | TGCGCACCCTTAGCGAGAGGTTTATCATTAAGGTCAACCTCTGGATGTTGTTTCGGCATCCTGCATTGAATCTGAGTTACTGTCTGTTTTCCTTGTTGGAAGGTTTGAAGAGCCAACCAGGAAACCCGTTTTTTCTGACGTAAGGGTGCGCA | A |
| Eco1 a1/a2 Length | 13 | 5281 | ATGCGCACCCTTAGCGAGAGGTTTATCATTAAGGTCAACCTCTGGATGTTGTTTCGGCATCCTGCATTGAATCTGAGTTACTGTCTGTTTTCCTTGTTGGAAATTGCGGGTAGCCAACCAGGAAACCCGTTTTTTCTGACGTAAGGGTGCGCAT | A |
| Eco1 a1/a2 Length | 14 | 5282 | TATGCGCACCCTTAGCGAGAGGTTTATCATTAAGGTCAACCTCTGGATGTTGTTTCGGCATCCTGCATTGAATCTGAGTTACTGTCTGTTTTCCTTGTTGGAAGAATGGAAAAGCCAACCAGGAAACCCGTTTTTTCTGACGTAAGGGTGCGCATA | A |
| Eco1 a1/a2 Length | 15 | 5283 | GTATGCGCACCCTTAGCGAGAGGTTTATCATTAAGGTCAACCTCTGGATGTTGTTTCGGCATCCTGCATTGAATCTGAGTTACTGTCTGTTTTCCTTGTTGGAATCGCCGAATAGCCAACCAGGAAACCCGTTTTTTCTGACGTAAGGGTGCGCATAC | A |
| Eco1 a1/a2 Length | 16 | 5284 | CGTATGCGCACCCTTAGCGAGAGGTTTATCATTAAGGTCAACCTCTGGATGTTGTTTCGGCATCCTGCATTGAATCTGAGTTACTGTCTGTTTTCCTTGTTGGAAGGGAGTTCCAGCCAACCAGGAAACCCGTTTTTTCTGACGTAAGGGTGCGCATACG | A |
| Eco1 a1/a2 Length | 17 | 5285 | CCGTATGCGCACCCTTAGCGAGAGGTTTATCATTAAGGTCAACCTCTGGATGTTGTTTCGGCATCCTGCATTGAATCTGAGTTACTGTCTGTTTTCCTTGTTGGAAAAATGGTACAGCCAACCAGGAAACCCGTTTTTTCTGACGTAAGGGTGCGCATACGG | A |
| Eco1 a1/a2 Length | 18 | 5286 | TCCGTATGCGCACCCTTAGCGAGAGGTTTATCATTAAGGTCAACCTCTGGATGTTGTTTCGGCATCCTGCATTGAATCTGAGTTACTGTCTGTTTTCCTTGTTGGAAGCTTCTGATAGCCAACCAGGAAACCCGTTTTTTCTGACGTAAGGGTGCGCATACGGA | A |
| Eco1 a1/a2 Length | 19 | 5287 | TTCCGTATGCGCACCCTTAGCGAGAGGTTTATCATTAAGGTCAACCTCTGGATGTTGTTTCGGCATCCTGCATTGAATCTGAGTTACTGTCTGTTTTCCTTGTTGGAATATATATAAAGCCAACCAGGAAACCCGTTTTTTCTGACGTAAGGGTGCGCATACGGAA | A |
| Eco1 a1/a2 Length | 20 | 5288 | ATTCCGTATGCGCACCCTTAGCGAGAGGTTTATCATTAAGGTCAACCTCTGGATGTTGTTTCGGCATCCTGCATTGAATCTGAGTTACTGTCTGTTTTCCTTGTTGGAAGGGTCGGGTAGCCAACCAGGAAACCCGTTTTTTCTGACGTAAGGGTGCGCATACGGAAT | A |
| Eco1 a1/a2 Length | 21 | 5289 | GATTCCGTATGCGCACCCTTAGCGAGAGGTTTATCATTAAGGTCAACCTCTGGATGTTGTTTCGGCATCCTGCATTGAATCTGAGTTACTGTCTGTTTTCCTTGTTGGAAGGAGATGTAAGCCAACCAGGAAACCCGTTTTTTCTGACGTAAGGGTGCGCATACGGAATC | A |
| Eco1 a1/a2 Length | 22 | 5290 | AGATTCCGTATGCGCACCCTTAGCGAGAGGTTTATCATTAAGGTCAACCTCTGGATGTTGTTTCGGCATCCTGCATTGAATCTGAGTTACTGTCTGTTTTCCTTGTTGGAAATTGGATGCAGCCAACCAGGAAACCCGTTTTTTCTGACGTAAGGGTGCGCATACGGAATCT | A |
| Eco1 a1/a2 Length | 23 | 5291 | AAGATTCCGTATGCGCACCCTTAGCGAGAGGTTTATCATTAAGGTCAACCTCTGGATGTTGTTTCGGCATCCTGCATTGAATCTGAGTTACTGTCTGTTTTCCTTGTTGGAATCAACTACGAGCCAACCAGGAAACCCGTTTTTTCTGACGTAAGGGTGCGCATACGGAATCTT | A |
| Eco1 a1/a2 Length | 24 | 5292 | TAAGATTCCGTATGCGCACCCTTAGCGAGAGGTTTATCATTAAGGTCAACCTCTGGATGTTGTTTCGGCATCCTGCATTGAATCTGAGTTACTGTCTGTTTTCCTTGTTGGAACCAAGTTCGAGCCAACCAGGAAACCCGTTTTTTCTGACGTAAGGGTGCGCATACGGAATCTTA | A |
| Eco1 a1/a2 Length | 25 | 5293 | ATAAGATTCCGTATGCGCACCCTTAGCGAGAGGTTTATCATTAAGGTCAACCTCTGGATGTTGTTTCGGCATCCTGCATTGAATCTGAGTTACTGTCTGTTTTCCTTGTTGGAAGAGTAGGTCAGCCAACCAGGAAACCCGTTTTTTCTGACGTAAGGGTGCGCATACGGAATCTTAT | A |
| Eco1 a1/a2 Length | 26 | 5294 | GATAAGATTCCGTATGCGCACCCTTAGCGAGAGGTTTATCATTAAGGTCAACCTCTGGATGTTGTTTCGGCATCCTGCATTGAATCTGAGTTACTGTCTGTTTTCCTTGTTGGAAGGGTCTCTTAGCCAACCAGGAAACCCGTTTTTTCTGACGTAAGGGTGCGCATACGGAATCTTATC | A |
| Eco1 a1/a2 Length | 27 | 5295 | TGATAAGATTCCGTATGCGCACCCTTAGCGAGAGGTTTATCATTAAGGTCAACCTCTGGATGTTGTTTCGGCATCCTGCATTGAATCTGAGTTACTGTCTGTTTTCCTTGTTGGAAAGTGCTTAAAGCCAACCAGGAAACCCGTTTTTTCTGACGTAAGGGTGCGCATACGGAATCTTATCA | A |
| Eco1 a1/a2 Length | 28 | 5296 | ATGATAAGATTCCGTATGCGCACCCTTAGCGAGAGGTTTATCATTAAGGTCAACCTCTGGATGTTGTTTCGGCATCCTGCATTGAATCTGAGTTACTGTCTGTTTTCCTTGTTGGAAGATCACCGGAGCCAACCAGGAAACCCGTTTTTTCTGACGTAAGGGTGCGCATACGGAATCTTATCAT | A |
